# Distant Ribose 2’-O-Methylation of 23S rRNA Helix 69 Pre-Orders the Capreomycin Drug Binding Pocket at the Ribosome Subunit Interface

**DOI:** 10.1101/2024.11.05.619916

**Authors:** Suparno Nandi, Debayan Dey, Pooja Srinivas, Christine M. Dunham, Graeme L. Conn

## Abstract

Loss of ribosomal RNA (rRNA) modifications incorporated by the intrinsic methyltransferase TlyA results in reduced sensitivity to tuberactinomycin antibiotics such as capreomycin. However, the mechanism by which rRNA methylation alters drug binding, particularly at the distant but functionally more important site in 23S rRNA Helix 69 (H69), is currently unknown. We determined high-resolution cryo-electron microscopy structures of the *Mycolicibacterium smegmatis* 70S ribosome with or without the two ribose 2’-O-methyl modifications incorporated by TlyA. In the unmodified ribosome, the tip of H69 adopts a more compact conformation, positioning two key nucleotides (A2137 and C2138) such that interactions with capreomycin would be lost and the binding pocket partially occluded. In contrast, methylation of 23S rRNA nucleotide C2144 results in conformational changes that propagate from the site of modification to the H69 tip, resulting in its movement away from h44, a more favorable positioning of C2138 and adoption of a more open conformation to enable capreomycin binding. Methylation of h44 also results in structural rearrangements at the H69-h44 interface that further support antibiotic binding. These structures thus reveal the effect and regulation of distant rRNA methylation on ribosome-targeting antibiotic binding.

## INTRODUCTION

Ribosome-targeting antibiotics have been critical to our ability to treat bacterial infections, but these essential drugs are now threatened by multiple resistance mechanisms (1). One major mode of resistance is conferred by antibiotic-resistance *S*-adenosyl-L-methionine (SAM)-dependent ribosomal RNA (rRNA) methyltransferases (2). For example, 23S rRNA methyltransferases of the Erm family are a clinically prevalent mechanism of resistance to macrolide antibiotics, while more recently, the aminoglycoside-resistance 16S rRNA methyltransferases have emerged as a new threat to even the most recently approved aminoglycoside antibiotics (3-5).

In contrast to these methyltransferases that confer antibiotic resistance by introducing rRNA modifications, loss of intrinsic rRNA methylation can also result in reduced antibiotic efficacy. For instance, deletion of the 16S rRNA methyltransferases KsgA (6) or RsmG (7) reduced susceptibility to the antibiotics kasugamycin and streptomycin, respectively. Similarly, loss of the cytidine-2’-O-methyltransferase TlyA results in decreased susceptibility to tuberactinomycin antibiotics such as capreomycin and viomycin (8). TlyA is encoded by all mycobacterial species, including the causative agent of tuberculosis (TB) *Mycobacterium tuberculosis (Mtb)*, and loss of the rRNA modifications incorporated by TlyA results in clinical resistance to capreomycin, a drug previously used for treating multidrug-resistant TB (9,10).

TlyA enzymes are divided into two subgroups, which incorporate either a single ribose modification in 23S rRNA of the large subunit (Type I) or two ribose modifications via dual 16S/23S rRNA activity (Type II). Mycobacterial species, including *Mtb* and *Mycolicibacterium smegmatis (Msm)*, encode Type II enzymes and thus have ribosomes modified with Cm2158^*Mtb*^/Cm2144^*Msm*^ and Cm1402^*Mtb*^/Cm1392^*Msm*^ on 23S rRNA Helix 69 (H69) and 16S rRNA helix 44 (h44), respectively. Note that hereafter, the terms methylation/modification are used to denote TlyA^II^-mediated dual methylation and *Msm* nucleotide numbering is used unless otherwise noted. Methylation on both nucleotides is essential for optimal capreomycin binding and its resulting antimicrobial activity (11). However, expression of *Thermus thermophilus (Tth)* Type I TlyA in *Escherichia coli* (*Eco*; which lacks TlyA) resulted in a 4-fold increase in sensitivity to capreomycin and viomycin, suggesting a greater relative contribution by the H69 modification despite its greater distance (∼21 Å) from the drug binding site (11). X-ray crystallographic and single-particle cryo-electron microscopic (cryo-EM) structures of the *Mtb* (12), *Tth* (13), and *Msm* (14) 70S ribosomes with capreomycin show that the antibiotic binds at the subunit interface created by H69 and h44, adjacent to the aminoacyl (A) site. Antibiotic binding at the H69-h44 interface has been proposed to alter the position of several H69 and h44 nucleotides and thus their interactions with the A-site tRNA, inhibiting either the decoding step and/or tRNA translocation (1,13,15).

While the necessity of rRNA methylation for optimal capreomycin binding to mycobacterial ribosomes is well documented, the molecular mechanism by which these modifications facilitate the drug-rRNA interaction is unclear. Methylated 16S rRNA C1392 is located at the drug-binding pocket, but as this residue appears to form only weak hydrogen bonds (>4 Å) (8,14,16), its modification cannot directly coordinate antibiotic binding. In structures of ribosomes from *Staphylococcus aureus* (17) and *Listeria innocua* (18) that lack TlyA and thus its associated methylations, the tip of H69 moves towards h44, adopting a more compact conformation with the H69 nucleobases oriented away from the drug binding pocket. In contrast, in TlyA^II^-modified *Msm* without capreomycin, the phosphate backbone of the H69 tip (nucleotides A2136-U2139) shifts away from h44 by ∼3 Å, repositioning the nucleobases of H69 to face the capreomycin binding pocket (14). Additionally, a structure of the TlyA^II^-bound *Msm* 50S subunit in a post-catalytic state, with C2144 modified with a SAM analog, captured the tip of H69 shifted away from h44 by ∼7 Å (19). From these observations, we and others (8,11) speculated that the more compact conformation of H69 somehow results from the lack of both modifications deposited by TlyA^II^, and results in greater occlusion of the capreomycin binding pocket. In contrast, H69 rRNA methylation by TlyA, and the altered H69 conformation that results, could favor interaction with capreomycin resulting in the observed rRNA modification-dependent sensitivity to the drug.

Here, we report high-resolution cryo-EM structures of the *Msm* 70S ribosome with and without the two rRNA modifications incorporated by TlyA^II^. These structures and complementary molecular dynamics studies reveal the critical role of C2144 methylation and propagation of its effect on the RNA helical structure of H69, including allosteric changes produced at the distant drug binding pocket at the tip of H69. In addition, our findings suggest a supporting role of C1392 methylation in maintaining the compact H69 structure and the overall architecture of the capreomycin binding pocket in the absence of the drug. RNA modifications that mediate allosteric changes in antibiotic binding sites, such as those revealed here for the tuberactinomycin antibiotics, thus represent an important new consideration for designing more effective small molecules targeting the bacterial ribosome.

## MATERIAL AND METHODS

### Purification of *Msm* 70S ribosomes

Ribosomes were isolated from *Msm* strains with (LR222) or without (LR222 C101A) TlyA^II^, as reported previously (20). Briefly, single colonies of *Msm* LR222 or LR222 C101A were used to inoculate 20 ml Middlebrook 7H9 liquid medium and grown overnight at 37 °C with mild agitation (shaking at 100 rpm). Fresh Middlebrook 7H9 medium (1-2 L) was inoculated with the overnight culture (1:100 dilution), and the cultures were grown at 37 °C with mild agitation over 35-41 hrs to an OD_600_ of ∼4.0. Cells were collected by centrifugation at 4,000 × *g* for 10 mins at 4 °C, washed twice (500 ml/L culture) with Buffer 1 (10 mM HEPES/ KOH pH 7.6, 1 M NH_4_Cl, 10 mM MgCl_2_, and 6 mM β-mercaptoethanol), and then once with the same buffer but with 0.1 M NH_4_Cl (Buffer 2).

Cells were resuspended in 5 ml of Buffer 2 per gram of wet cell mass and lysed by two passes through an EmulsiFlex-C5 high-pressure homogenizer (Avestin). After adding DNase I (10 U/mL lysate), the lysate was cleared by centrifugation for 10 and 20 mins at 19,400 × *g* and 27,000 × *g*, respectively. Ribosomes were pelleted by ultra-high-speed centrifugation of the supernatant at 278,800 × *g* for 18 hrs. To remove bound tRNA, ribosome subunits were split by resuspension and dialysis against Buffer 3 (10 mM HEPES/KOH pH 7.6, 0.1 M NH_4_Cl, 0.3 mM MgCl_2_, and 6 mM β-mercaptoethanol), followed by reassociation via subsequent overnight dialysis against Buffer 2. Reassembled 70S ribosomes were separated from individual subunits by centrifugation (90,200 × *g*) on a 10 to 40% sucrose gradient for 18 hrs at 4 °C, and subsequent fractionation using an ÄKTA Purifier10 system. Pooled 70S fractions were centrifuged at 226,000 × *g* for 18 hrs at 4 °C and the pelleted 70S resuspended, dialyzed overnight against Buffer 2, and flash-frozen for storage at -80 °C as 20 µl aliquots of LR222 (2.55 µM) and LR222 C101A (2.39 µM).

### Cryo-EM specimen preparation, data collection, and structure determination

Ribosomes were diluted to 1.0 and 3.0 µM for LR222 (TlyA-methylated) and LR222 C101A, respectively. Each sample (3 µl) was applied to glow-discharged Quantifoil Cu R1.2/1.3 300 mesh grids, blotted for 4 s, and, after a 10 s wait time, vitrified in liquid ethane using a Vitrobot Mark IV (Thermo Fisher Scientific) operating at room temperature and 100% relative humidity.

Cryo-EM micrographs (5060 and 5665 for LR222 and LR222 C101A, respectively) were recorded as movies using Leginon (21) with defocus range of -0.6 to -2.2 µm and -0.6 to -2.1 µm at 81,000× magnification (1.069 Å/pixel) on a Titan Krios 300 kV microscope with Gatan K3 direct electron detector at the National Center for CryoEM Access and Training (NCCAT). The total dose was 52.51 e^-^ /Å^2^, distributed over 40 frames for 2.0 s (50 ms per frame), resulting in a dose rate of 1.31e^-^/Å per frame.

The complete workflow for cryo-EM structure determination is summarized in **Supplementary Figs. S1** and **S2**. First, aligned and dose-weighted images were imported, and the contrast transfer function (CTF) estimate was accomplished using Gctf in Relion-3.1 (22,23). Automated particle picking was guided by an initial Laplacian-of-Gaussian-based autopicking of 200 images, which resulted in 35,443 and 42,841 particles for LR222 and LR222 C101A, respectively. Two-dimensional (2D) classes of these particles were used as a reference for subsequent autopicking with each complete image data set, resulting in the selection of 3,878,842 and 4,579,196 particles (400-pixel box size, 1.069 Å/pixel), respectively, which were 4×4 binned (100-pixel box size, 4.276 Å/pixel). Following the 2D classification of each particle set, an initial map was created from the classes and used to generate 3D maps by iterative rounds of 3D refinement and 3D classification to remove non-ribosomal particles. The particles corresponding to the final 3D classes for both reconstructions were then unbinned at full resolution (400-pixel box size, 1.069 Å/pixel), and following a final round of 3D classification to remove unsuitable classes for the non-TlyA methylated ribosome, multiple rounds of 3D and CTF refinements resulted in post-processed maps of 3.17 Å and 3.24 Å for LR222 (TlyA methylated) and LR222 C101A (non-methylated) ribosomes, respectively.

To improve the resolution of the maps, multibody refinement (24) was also performed on the final selected particles with individual masks corresponding to the large and small subunits of each ribosome. This process resulted in four maps with resolutions of 3.08 Å (50S) and 3.19 Å (30S) for LR222, and 3.17 Å (50S) and 3.34 Å (30S) for LR222 C101A. Postprocessing of these maps without sharpening did not result in further enhancement of resolution based on gold-standard refinement Fourier Shell Correlation (0.143 cutoff) (**Supplementary Fig. S3A-C,G-I**). The postprocessed maps were subsequently sharpened using PHENIX (adjusted surface area) (25). Additionally, DeepEMhancer (Sanchez-Garcia et al., 2021) was used to sharpen the half-maps obtained from multibody refinement of the individual subunits in both methylated and unmethylated ribosomes. Relion-3.1 was used to generate local resolution maps (**Supplementary Fig. S3D-F,J-L**).

The 3D refined map was used to build the initial model for both the methylated and unmethylated ribosomes. An *Msm* 70S structure (PDB code 5ZEB) without tRNA was docked in the map for each ribosome and refined using global minimization, rigid body, local grid search, simulated annealing, and B factor refinement in PHENIX (26). Subsequently, each 70S model was split into individual 50S and 30S subunits, and each docked in its corresponding multibody refined map using Chimera (v1.16) (27). Each subunit model was further refined using the same strategies in PHENIX. Finally, the individual subunit models were merged to create a final complete model of 70S followed by final model adjustment in COOT (v0.9.8.1) using the DeepEMhancer sharpened multibody refinement maps for each subunit (28). Both structures were validated using PHENIX, and the final parameters for data collection, processing, model construction, refinement, and validation are provided in **Table 1**. Structural images were prepared using PyMOL (v2.5.4) (29) or ChimeraX (v1.7) (30) using the DeepEMhancer sharpened multibody refinement maps for each subunit.

**Table 1:**
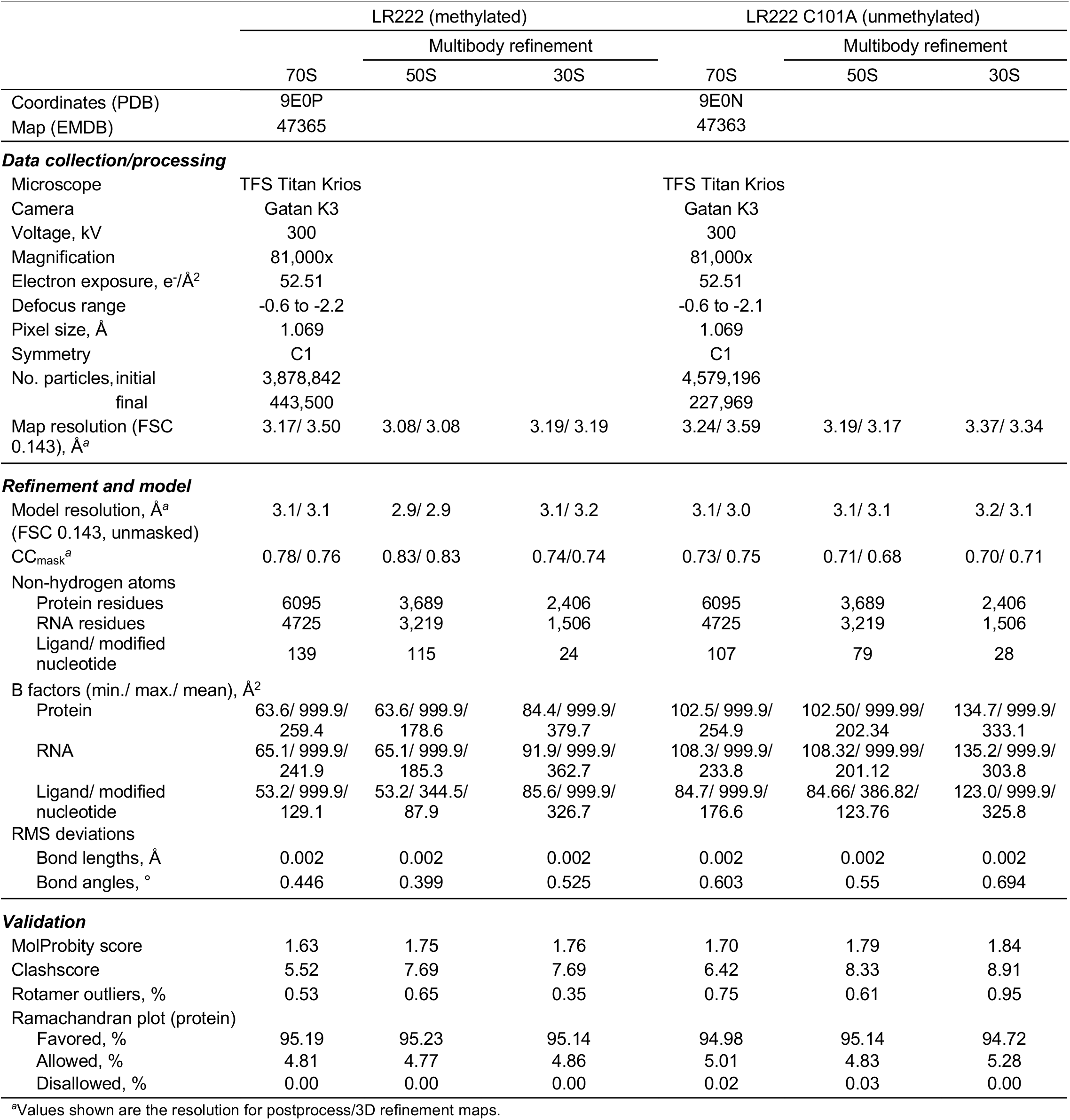
Cryo-EM data collection, refinement, and model validation for the methylated and unmethylated *M. smegmatis* 70S ribosome.

### Molecular dynamics (MD) simulations

The 19-nucleotide segment of 23S rRNA H69 (nucleotides 2129-2148), with and without the C2144 ribose methylation, was used for partially restrained classical MD simulations using Desmond and the OPLS4 force field from the Schrödinger software (2023-4), which accommodates modified RNA residues. One thing to note is that H69 does not interact with any proteins and thus the MD simulations only included RNA. Restraints (spring constant, k = 1) were applied to the terminal nucleotides (C2129-G2131 and U2147-C2148) to maintain structural stability. Since the terminal residues in the RNA fragment are part of rRNA, these restraints were applied to prevent the RNA from unwinding or opening unrealistically at the terminal regions. Prior to simulation, each system was neutralized by adding Na^+^ ions around the H69 RNA in the Schrödinger software System Builder module. The neutralized RNA was then immersed in TIP3P water, and random water molecules were replaced with Na^+^ ions to achieve a total ionic strength of 150 mM NaCl. The solvated system underwent relaxation and equilibration in the isobaric–isothermal (NPT) ensemble (P = 1 atm, T = 310 K) for 10 ns. Subsequently, production simulations (100 ns) were conducted using the Nose-Hoover chain and Martyna-Tobias-Klein thermostat and barostat with relaxation times of 1 and 2 ps, respectively, in the canonical (NPT) ensemble using the last configuration from the equilibration. The equations of motion were integrated using multiple time steps for short-range (2 fs) and long-range (6 fs) interactions, with a 9-Å cutoff for nonbonded interactions. Coordinates were saved every 100 ps. Three replicates of these simulations were performed. Root mean square deviation (RMSD)-based clustering was performed on each trajectory with 10-frame intervals on all non-hydrogen (heavy) atoms. The top 5-7 frames, covering over 60% of all conformations, were selected for analysis of intrabase and interbase pair helical parameters.

### Calculation of H69 helical (base step) parameters

The RNA helical parameters for H69 of LR222 (TlyA methylated) and LR222 C101A (unmethylated) ribosomes extracted from the 70S structures or from MD simulations were calculated using the Web 3DNA 2.0 server (31). Values for selected parameters were plotted in GraphPad Prism (v9.5.1.733).

## RESULTS

### Structures of TlyA^II^-methylated and unmethylated *Msm* 70S ribosomes

*Msm* methylated (i.e, with TlyA^II^; strain LR222) and unmethylated (i.e., without TlyA^II^; strain LR222 C101A) 70S ribosomes were purified under identical conditions as described previously (20), and their structures determined by single-particle cryo-EM at overall resolutions of 3.16 Å and 3.24 Å, respectively (**Fig. 1**). Multibody refinement resulted in similar overall resolution maps for the 50S and 30S subunit regions with slight improvement for the 50S subunit: methylated 50S (3.08 Å) and 30S (3.19 Å) and unmethylated 50S (3.17 Å) and 30S (3.37 Å) subunits. More importantly, multibody refinement improved the map quality for the region corresponding to H69, with local resolution analysis revealing similar values for H69 as the majority of the corresponding full 50S subunit (**Supplementary Fig. S3E,K**). The maps from multibody refinement were subsequently postprocessed without sharpening and sharpened using PHENIX (adjusted surface area) (25) and DeepEMhancer (32), and the final model was generated by PHENIX-based structural refinement (**Supplementary Figs. S1 and S2**).

**Figure 1.**
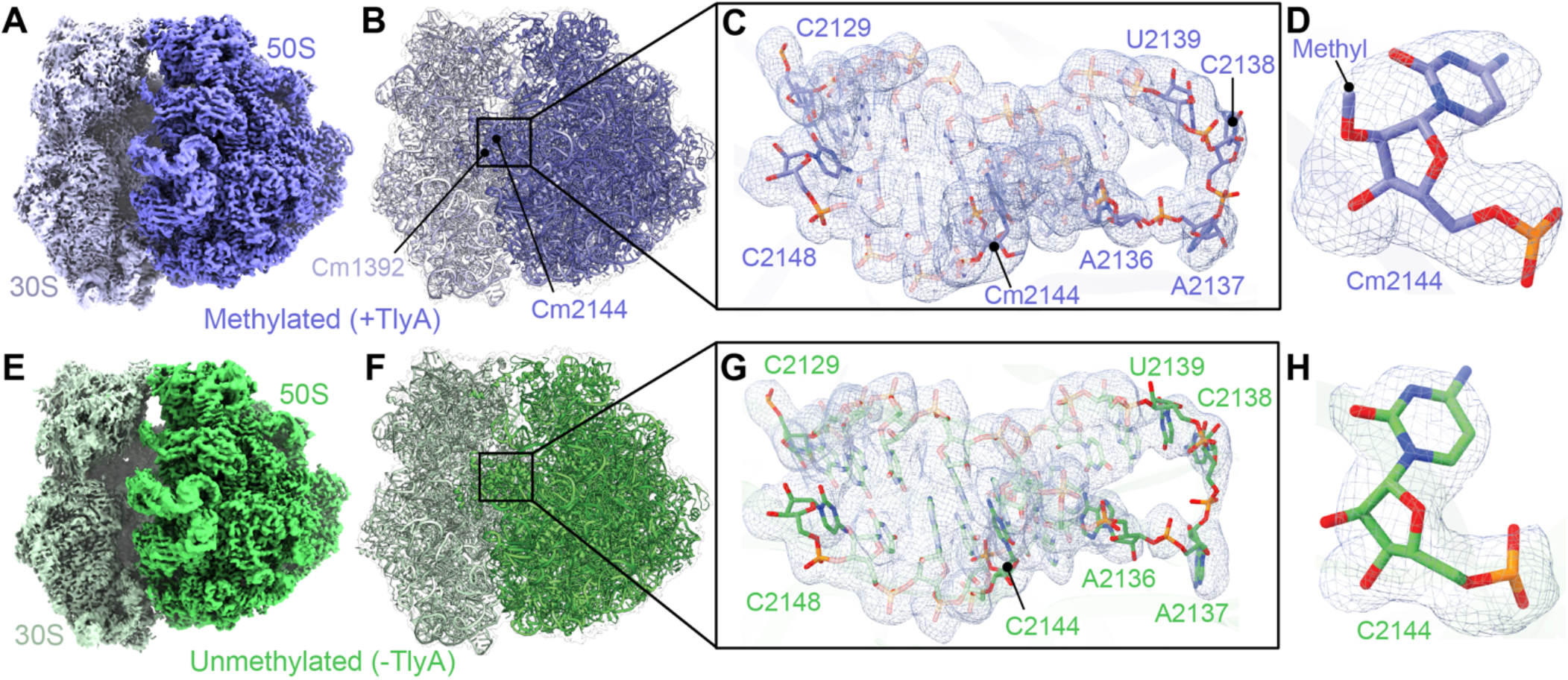
Cryo-EM map and model of the TlyA-methylated and unmethylated H69 within the *Msm* 70S ribosome. ***A***, Postprocessed (PHENIX sharpened) map of the methylated 70S ribosome (50S and 30S subunits shown in blue and pale blue, respectively) at 3.16 Å resolution. ***B***, The final model shown within a semi-transparent white map and highlighting the location of H69 (boxed region) and indicating the sites of methylation on both subunits. ***C***, Zoomed-in view of the indicated region, showing the H69 structure with a surrounding map from 50S multibody refinement (DeepEMhancer sharpened, map threshold 0.01; map value range –0.00171 to 1.86). ***D***, Methylated Cm2144 shown within the map (DeepEMhancer sharpened, map threshold 0.04; map value range – 0.00171 to 1.86) supporting the presence of the expected 2’-O-methyl group. ***E-H***, As for *panels A-D* but for the unmethylated *Msm* 70S ribosome (50S and 30S subunits in green and light green, respectively) at 3.24 Å resolution, with the unmodified H69 (C2144) highlighted (boxed region). For *panels G* and *H*, the map is DeepEMhancer sharpened with a map threshold of 0.013 and 0.042, respectively; map value range –0.00235 to 2.04. For panels showing stick models in this and other figures, O, N, P, and C atoms are shown in red, blue, orange, and backbone color, respectively.

This process and the resulting maps enabled modeling of the entire H69, including its terminal tip residues, A2136-U2139, which form a four nucleotide RNA loop and comprise part of the capreomycin binding pocket (**Fig. 1C,G**). The modification of C2144 in the methylated, but not unmethylated, 70S ribosome could be visualized, and the 2’-O methylated nucleotide confidently modeled (**Fig. 1D,H**). Finally, the separate 50S and 30S models were combined to create complete structures of the methylated and unmethylated 70S ribosomes (**Fig. 1** and **Supplementary Figs. S1** and **S2**). Overall, the methylated and unmethylated ribosome structures are essentially identical, with a RMSD of ∼0.6 Å over 4652 phosphate backbone atoms. Additionally, both structures show the expected mycobacterial ribosome-specific protein and 23S rRNA expansions (14), although the map quality at the tip of the extended H54a helix was fragmented in both reconstructions. Consistent with an earlier structure (19), 23S rRNA H54a in the unmethylated *Msm* 70S ribosome is flexible as the tip of the helix was observed to be shifted by ∼54 Å compared to another unmethylated *Msm* 50S ribosome (PDB code7S0S) even though their overall structures are again essentially identical (RMSD ∼0.7 Å). Thus, our structures of TlyA^II^-methylated (LR222) and unmethylated (LR222 C101A) ribosomes appear virtually identical to one another and prior structures, indicating that the modifications incorporated by TlyA^II^ do not induce large-scale changes in 70S ribosome.

### C2144 2’-O-methylation results in a more open conformation of the tip of H69

Alignment of H69 from our methylated *Msm* 70S ribosome with that of a previously published *Msm* 70S structure (PDB code 5O61) reveals that these two H69 structures are very similar **Supplementary Fig. S4A** and **Table S1**). Likewise, a comparison of our unmethylated *Msm* ribosome and those from two bacteria lacking any TlyA, *L. innocua* (PDB code 8UU4), and *S. aureus* (PDB code 7TTU), reveals that H69 is structurally conserved among unmethylated ribosomes (**Supplementary Fig. S4B** and **Table S1**). Direct comparison of H69 between our two new structures, with and without methylation by TlyA^II^, reveals that the only significant difference is at the apical tip of the H69 helix (nucleotides A2136-U2139; **Fig. 2A** and **Table S1**). In the methylated H69, the phosphodiester backbone shifts by ∼3 Å to create a more open conformation of these H69 loop nucleotides, whereas the corresponding region of the unmethylated ribosome adopts a more compact structure.

**Figure 2.**
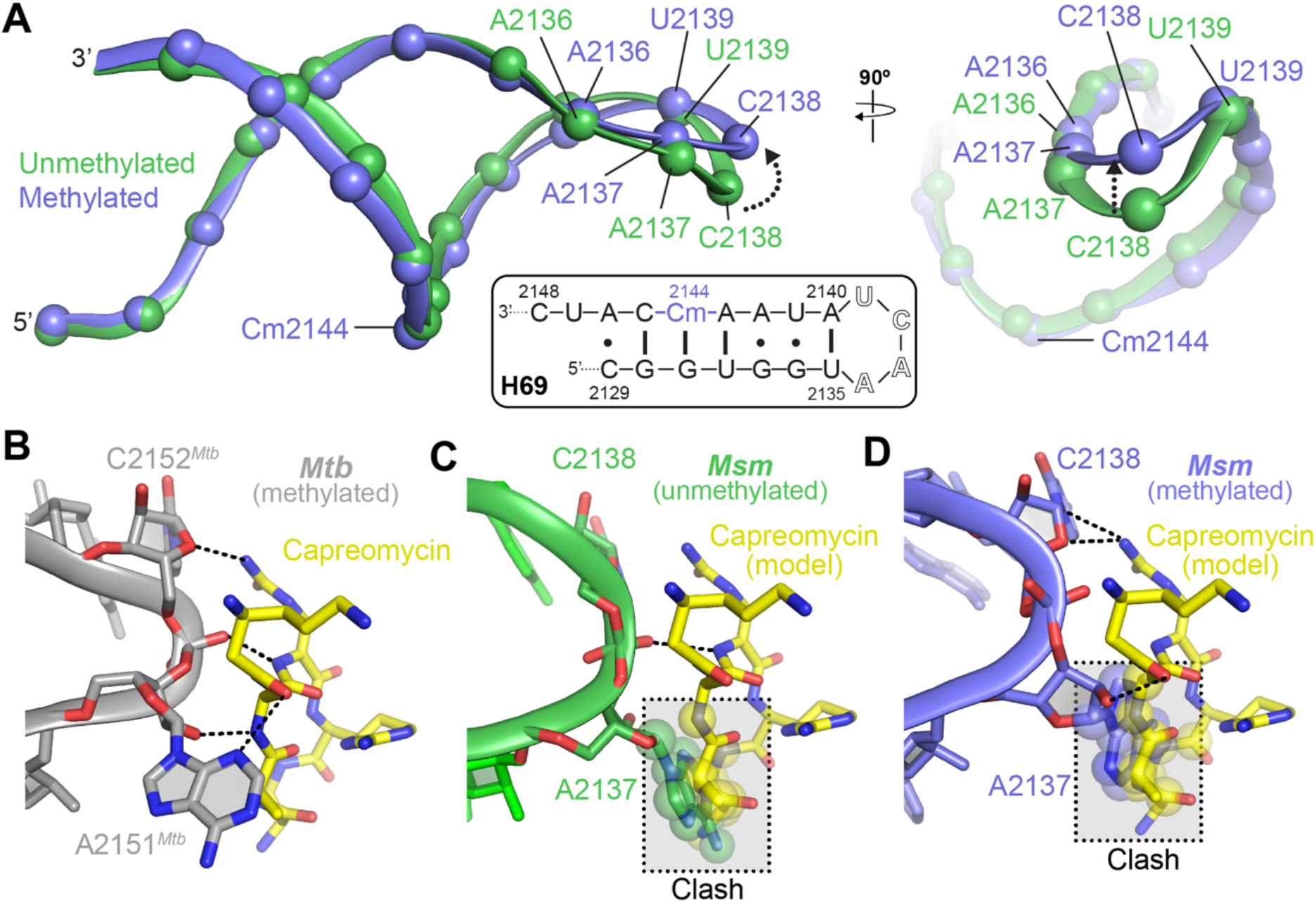
The nucleotides at the tip of H69 undergo changes in the methylated vs. unmethylated ribosomes, potentially impacting capreomycin binding. ***A***, Two orthogonal views of a superimposition of the methylated (blue) and unmethylated (green) H69 shown backbone and phosphate spheres. Differences in the tip are indicated with a dotted line arrow. *Inset*, shows the sequence and secondary structure of the 23S rRNA H69 region. ***B***, Residues A2151 and C2152 at the tip of H69 in *Mtb* (gray; PDB code 5V93) stabilize capreomycin (yellow) in its binding pocket via hydrogen bonding interactions (dashed lines). ***C***, In the unmethylated *Msm* ribosome, A2137 is predicted to clash (semi-transparent boxes) with the modeled capreomycin, while C2138 is no longer positioned to interact with the antibiotic. ***D***, In the methylated *Msm* ribosome, movement of C2138 and the tip of H69 partially opens the capreomycin binding pocket and allows the nucleotide to contact the modeled antibiotic. A2137 remains in a position predicted to result in clashes with the bound capreomycin and would thus require further movement promoted by drug binding.

Nucleotides A2137 (A2151^*Mtb*^/A1913^*Tth*^) and C2138 (C2152^*Mtb*^/C1914^*Tth*^) at the tip of H69 have been reported to help anchor capreomycin in its binding pocket at the subunit interface (12,13). Therefore, to assess the potential effect of the structural differences observed in our structures at the tip of H69 on antibiotic binding, we compared our two structures with the *Mtb* and *Tth* 70S ribosome-capreomycin complexes. Interestingly, all of H69 in these two drug-bound structures, including the phosphodiester backbone at the tip of the helix, is essentially identical to the unmethylated *Msm* 70S ribosome structure but with distinct orientations of the A2137 and C2138 nucleobases at the tip of H69 (**Supplementary Fig. S4C,D**), further supporting their role in positioning capreomycin (**Fig. 2B** and **Supplementary Fig. S5A**).

Next, to compare our structures with the capreomycin-bound *Mtb* and *Tth* H69, we modeled the antibiotic in the drug-binding pocket within our methylated and unmethylated H69. The distinct orientations of A2137 and C2138 in the unmethylated *Msm* 70S H69 suggest that C2138 would make either no interactions or a single potential hydrogen bond with the modeled capreomycin from *Tth* and *Mtb*, respectively, while A2137 is positioned in a manner that would occlude drug binding in both cases (**Fig. 2C** and **Supplementary Figs. S5B** and **S6A**). However, methylation of C2144, ∼21 Å away from these nucleotides in H69, results in a reorientation of C2138 to a position more similar to C2152^*Mtb*^/C1914^*Tth*^ thus potentially restoring favorable interactions with capreomycin, including putative hydrogen bonds made by both the ribose and nucleobase of C2138 (**Fig. 2D** and **Supplementary Figs. S5C** and **S6D**). These structures can thereby explain why the unmethylated H69 is less suitable for binding capreomycin, due to the loss of interactions made by C2138, resulting in the observed reduction in drug efficacy. In contrast, Cm2144 methylation positions C2138 to engage with capreomycin, which can support additional necessary reorganization (e.g., displacement of A2137) for full accommodation of the drug into its binding pocket.

### Allosteric signal propagation from Cm2144 results in a conformational change at the tip of H69

The site of methylation at C2144 is five nucleotides in the 3’ direction from U2139 (∼23 Å distant), the final base of the four-nucleotide loop at the tip of H69. To understand how this distant methylation might influence the capreomycin binding pocket conformation, we analyzed RNA helical parameters for H69 nucleotides C2129-C2148, including the twist, roll, tilt, rise, slide, and shift of each nucleotide pair (i.e., base steps), which previous studies have shown to be useful in defining allosteric signaling in DNA (33,34). Comparison of the methylated (Cm2144) and unmethylated (C2144) H69 structures revealed differences in each of these helical parameters for the region comprising nucleotides U2141 to A2136, i.e., base steps 8-12 between the site of methylation at C2144 and A2136, the first 5’ nucleotide of the H69 loop (**Fig. 3**).

**Figure 3.**
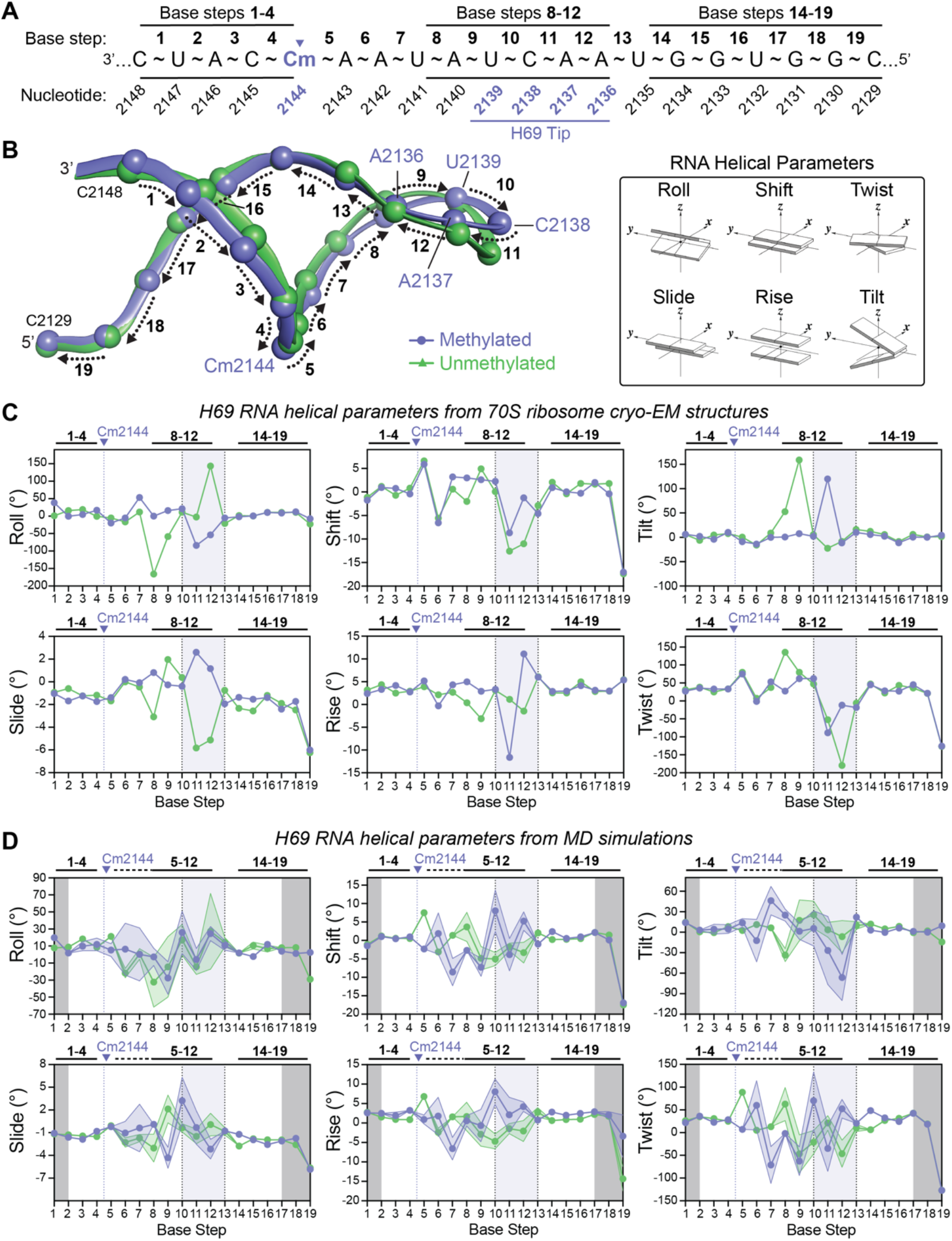
C2144 methylation results in propagation of helical conformational changes to the tip of H69. ***A***, Schematic of the H69 sequence shown in the direction of signal propagation (3’ to 5’) with nucleotide numbers and base steps from helical analysis in 3DNA shown below and above the sequence, respectively. Regions of the sequence noted in the text are indicated (*top*). ***B***, Superposition of the methylated (blue) and unmethylated (green) H69 structures with base step numbers indicated. ***C***, Comparison of the indicated H69 helical parameters (*inset*, above right) for methylated and unmethylated 70S ribosomes The dotted vertical line and shaded (grey) region indicate the site of methylation (Cm2144) and H69 tip, respectively. ***D***, Comparison of average values and standard deviation (color-coded shaded regions) for the same six helical parameters derived from representative structures from MD simulations of the methylated and unmethylated H69 structures. Light and dark grey shading denote the H69 tip and regions are those for which a restraint was applied during the simulation, respectively.

In contrast, no differences were observed in the other regions of the helix, i.e., for nucleotides U2135-C2129 of the complementary strand on the 5’ side of H69 (base steps 14-19) and nucleotides C2148-Cm2144 to the 3’ side of C2144 (base steps 1-4). We additionally compared the standard deviation for differences in each helical parameter averaged within each of these three distinct regions of H69. For all parameters assessed, these values were substantially higher for the region spanning from near the methylation site to U2139 (base steps 5-9), consistent with this region being the most impacted by the presence or absence of C2144 methylation (**Table S2**). Finally, helical parameters were calculated for H69 from other structures of methylated *Msm* (PDB code 5O61) and unmethylated *S. aureus* (PDB code 7TTU) ribosomes and these were also found to be consistent with our methylated and unmethylated structures, respectively, with only minor differences primarily localized at the tip of the helix for the unmethylated structure (**Supplementary Fig. S7A,B**). These observations further support the idea that methylation triggers alterations in helical structure, ultimately facilitating allosteric signal transmission that propagates in the 3’-direction from C2144 to the H69 tip, resulting in the observed shifts in the loop nucleotide positions.

To further investigate how C2144 ribose methylation impacts H69 conformational dynamics, we performed MD simulations of H69 nucleotides C2129 to C2148 with and without the modification. To prevent unrealistic unpairing in this isolated model RNA system, nucleotides C2129-G2131 (base steps 18-19) and U2147-C2148 (base step 1) at the base of the H69 stem, which are normally connected to adjacent rRNA nucleotides in the 70S ribosome, were partially restrained during the simulations (see **Materials & Methods**). Multiple conformations (five or more) of H69 from each simulation were selected using RMSD-based clustering of the trajectories, such that each set collectively represented >60% of all RNA conformations in the simulation. RNA helical parameters of these representative conformations were calculated, and the same six parameters compared as before (**Fig. 3D**). Again, significant differences between methylated and unmethylated H69 were observed between the site of methylation and the nucleotides of the loop at the tip of H69 (base steps 5-12). Additionally, the H69 regions to both the 5’- (base steps 13-17) and 3’-side (base steps 1-4) remain essentially identical between the methylated and unmethylated ribosomes and exhibit significantly less variability between the representative structures from each simulation (indicated by the shaded region on each plot in **Fig. 3D**). Thus, MD simulations further support the idea that methylation at C2144 imparts conformational and nucleotide dynamic changes on H69 that propagate exclusively from the site of modification back (3’ direction) to the tip of H69 where these changes manifest as allosteric alterations in the conformation of the capreomycin binding pocket.

### 16S rRNA C1392 methylation alters the conformation of the capreomycin binding pocket

Previously published structures of methylated *Mtb* and *Tth* 70S ribosomes with capreomycin showed that 16S rRNA nucleotide G1484^*Mtb*^/G1491^*Tth*^ (G1475) forms part of the drug binding site and directly interacts with the antibiotic (12,13). We asked whether the TlyA^II^ methylations play a role in supporting the formation of this interaction with capreomycin but found no change in the position of G1475 position which is similar in both our methylated and unmethylated structures, and in the drug-bound structures (**Supplementary Fig. S8**). Thus, G1475 appears to be oriented to interact with the antibiotic regardless of the methylation status of 23S rRNA C2144 and 16S rRNA C1392.

16S rRNA nucleotides A1391 (A1401^*Mtb*^/A1408^*Tth*^) and the TlyA-target 16S rRNA nucleotide C1392 (C1402^*Mtb*^/C1409^*Tth*^) have also been proposed to be important for capreomycin binding as they are positioned to make potential weak hydrogen bonding interactions with the bound drug at the subunit interface (13,16). However, as these potential hydrogen-bonding interactions distances exceed ∼4.0 Å (**Supplementary Fig. S9A,B**), we asked whether these two 16S rRNA nucleotides might have an alternative role in supporting the formation of the capreomycin binding pocket. The 23S and 16S rRNA subunit interface in this region of our methylated and unmethylated ribosomes align with an RMSD of 0.86 Å, indicating that they are overall structurally very similar (**Supplementary Fig. S9C**); however, local differences in H69 tip nucleotide positions are observed. In the unmethylated ribosome, C2138 at the tip of H69 interacts with C1392 of h44 (**Fig. 4A**, *left*, and **Supplementary Fig. S6B**), whereas in the methylated ribosome, since H69 becomes more compact, C2138 packs further into its helix and thereby moves away from C1392, resulting in a loss of these interactions (**Fig. 4A**, *center*, and **Supplementary Fig. S6E**). Notably, without these reorientations, the methyl group of C1392 would clash with the base of C2138 (**Fig. 4A**, *right*). As such, the methylation of C1392 may directly contribute to maintaining the more favorable position of C2138 for interaction with capreomycin. In contrast, A1391 is not positioned to form hydrogen bonds with A2137 in either the unmethylated or methylated ribosome, although the two nucleotides are closer in the methylated ribosome as compared to the unmethylated ribosome (**Fig. 4B**, *left* and **Supplementary Fig. S6B, E**). However, A1391 in the context of TlyA^II^-methylated h44 is slightly shifted towards A2137, while A2137 itself moves away from A1391, which ensures that potential steric clashes are avoided (**Fig. 4B**).

**Figure 4.**
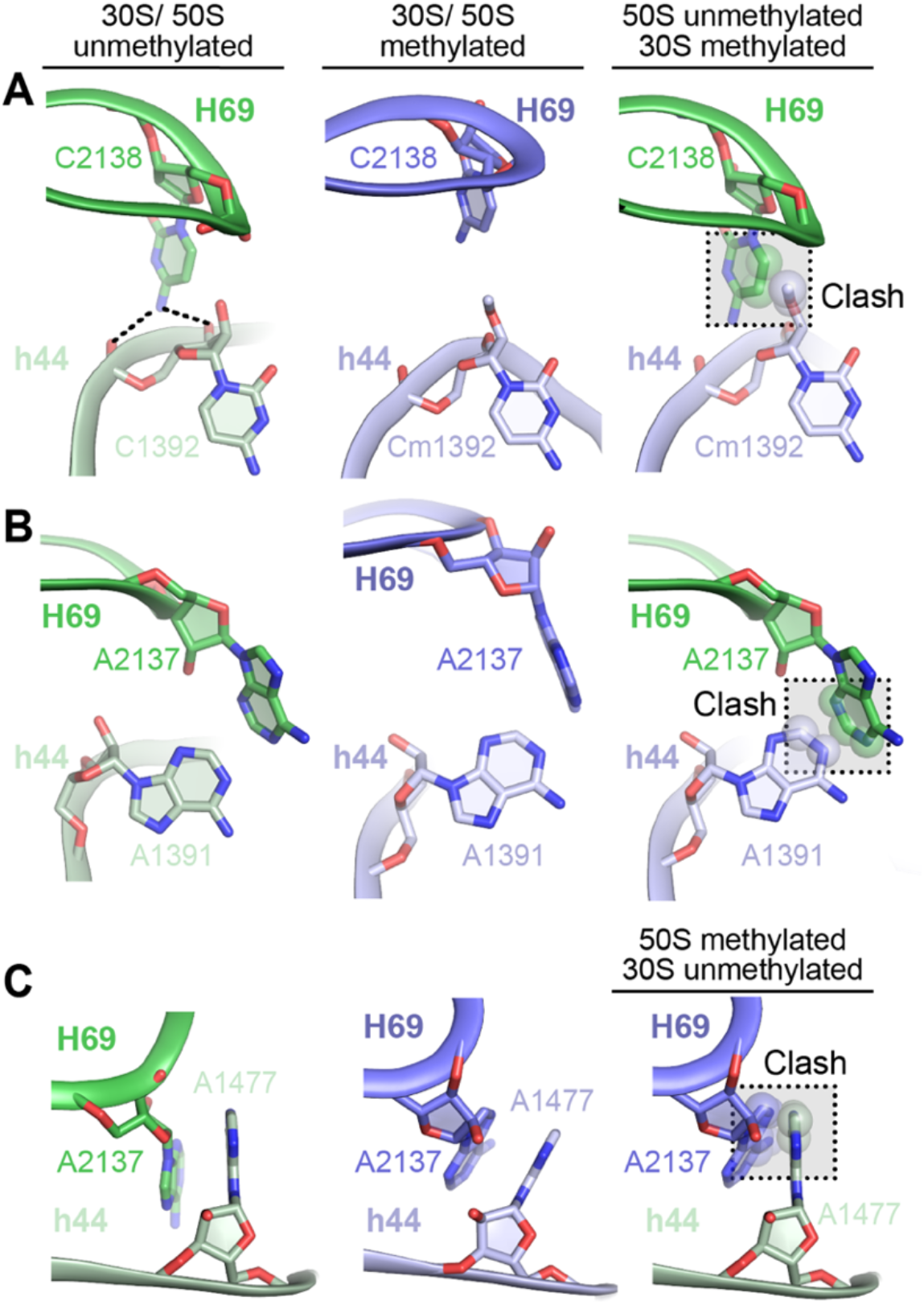
16S rRNA nucleotides assist in preserving the architecture of the capreomycin binding site. ***A***, In the unmethylated ribosome (*left*), 23S rRNA C2138 interacts with 16S rRNA C1392, while in the methylated ribosome (*center*), these contacts are lost due to the movement of C2138 away from Cm1392. A model combining C2138 as positioned in the unmethylated ribosome and Cm1392 (*right*) show that methylation would result in steric clashes (spheres) without this movement of C2138. Methylation of C1392 thus supports the maintenance of C2138 in the more open conformation of the H69 tip. In this and other panels, 16S and 23S rRNA nucleotides are shown in light and dark green (unmethylated) and blue (methylated), respectively. ***B***, 16S rRNA A1391 and 23S rRNA A2137 are adjacent in the unmethylated ribosome but do not directly interact (*left*). In the methylated 70S ribosome (*center*), A2137 moves away from A1391 as the tip of H69 moves upward. A model combining A1391 in the methylated 30S reveals potential clashes with A2137 when positioned as observed in the more compact H69 conformation of the unmethylated 50S subunit (*right*). ***C***, In the unmethylated ribosome (*left*), 23S rRNA A2137 and 16S rRNA A1477 make a π-stacking interaction that is is maintained by coordinated rotation of both nucleotides in the methylated ribosome (*center*). A model combining methylated 30S and unmethylated 50S subunits (*right*) shows steric clashes would result without the rotations, indicating that these movements maintain the stacking interaction and the more open conformation of H69 in the methylated ribosome.

Finally, another significant movement was observed for the h44 strand complementary to the strand containing A1391 and C1392. In the unmethylated ribosome, the nucleobase of A1477 of h44 makes a *π*-stacking interaction with the nucleobase of A2137 in the H69 tip (**Fig. 4C**, *left*, and **Supplementary Fig. S6A,C**). This contact is preserved in the methylated ribosome through rotations of A2137 and A1477, the absence of which would result in steric clash (**Fig. 4C** and **Supplementary Fig. S6D,F**). Previously published ribosome structures from different organisms, including *Eco* (35) and *Tth* (36,37) also show the presence of this contact, indicating its conservation in different species.

Therefore, our structures suggest that the methylation of h44 alters the positions and interactions of C1392, A1391, and A1477 in a manner that supports the changes at the tip of H69, allosterically mediated by C2144 methylation, that is necessary for optimal capreomycin binding and antimicrobial activity.

## DISCUSSION

We determined two cryo-EM structures of the *Msm* 70S ribosome that are identical except for the presence or absence of the two cytidine-2’-O-methylations incorporated by the methyltransferase TlyA^II^. Although the structures are highly similar overall, they reveal key conformational differences at the tip of H69 of the 50S subunit and reorganization of key residues in h44 of the 30S subunit that depend on the presence or absence of the two modifications. This work thus reveals how rRNA methylation can promote ribosome-targeting drug activity, in contrast to the more wide-spread mechanism of blocking antibiotic binding by steric clashes that then lead to resistance (2,38,39). In particular, we find that a 23S rRNA methylation of C2144 causes conformational changes at the distant capreomycin drug binding site as well as a reorganization of 16S rRNA nucleotides and a potential Cm1392 methylation-induced steric clash, all contribute to facilitating rather than impeding antibiotic binding. These changes appear to collectively underpin the established rRNA methylation-dependent activity of tuberactinomycin antibiotics such as capreomycin.

Nucleotides A2137 and C2138 at the tip of H69 adopt distinct positions in previously published structures of methylated *Mtb* or *Tth* 70S with bound capreomycin (12,13) as well as in the methylated and unmethylated *Msm* 70S structures presented here. In the unmethylated *Msm* ribosome, the tip of H69 adopts a more compact conformation, and is positioned close to h44 with A2137 and C2138 occupying the capreomycin binding pocket, presenting a steric hindrance to its binding (**Fig. 5A**). In the TlyA^II^-methylated ribosome, the tip of H69 adopts a more open conformation in which C2138 moves away from h44 as compared to the unmethylated ribosome; this reconfiguration opens up the capreomycin binding pocket in the fully assembled ribosome and positions C2138 to anchor the incoming antibiotic (**Fig. 5A**). Therefore, the previously reported elevated capreomycin minimal inhibitory concentration (MIC) against *Eco* (which lacks TlyA) (11) and the appearance of clinical capreomycin-resistant isolates with *tlyA* mutations that render the enzyme inactive (40,41) may be due to the lack of interactions between C2138 and the modeled capreomycin. TlyA-mediated methylation of C2144 brings C2138 closer to the antibiotic, promoting the drug-ribosome interaction and resulting in increased efficacy and the observed reduction in capreomycin MIC (11). Interestingly, even in the methylated *Msm* 70S ribosome structure, A2137 remains in a position that partially blocks the antibiotic binding site, whereas the structure of the *Tth* 70S ribosome-capreomycin complex shows a flipped orientation of A1913^*Tth*^ (A2137) that enables interactions between capreomycin and the A-site tRNA (13). Thus, the need for capreomycin binding to trigger movement of A2137 from its position in the drug-free binding pocket may underpin the mechanism of drug action by promoting this nucleotide’s interaction with the A-site tRNA, resulting in decoding defects and/or inhibition of translocation (15,42) (**Fig. 5A**).

**Figure 5.**
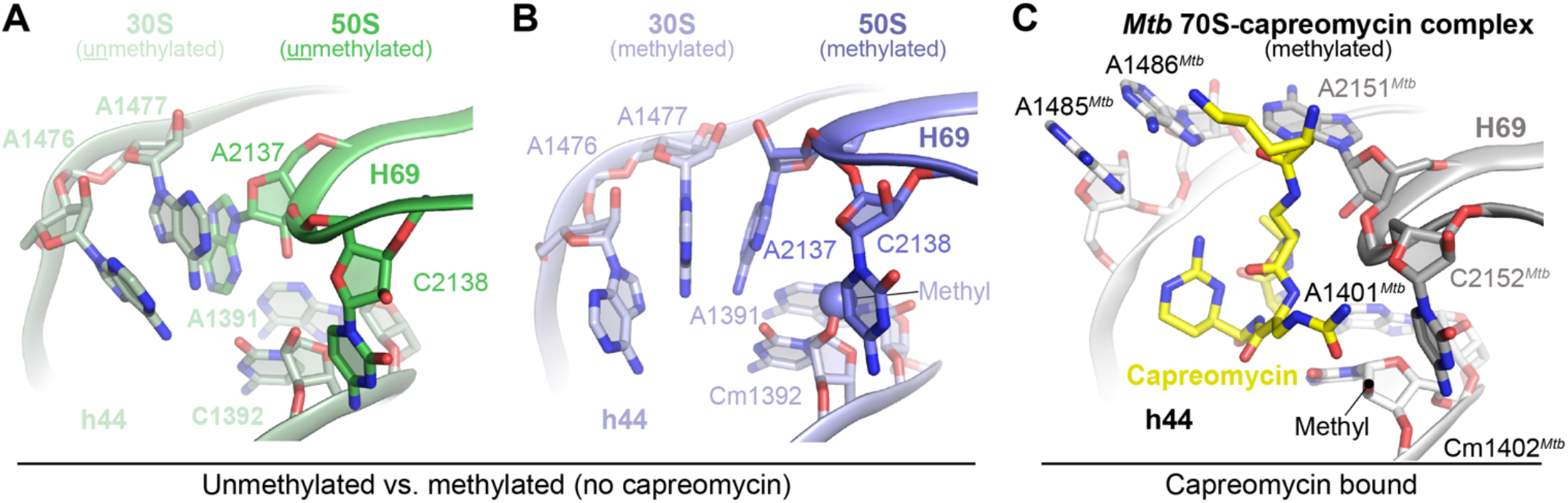
The capreomycin binding pocket is initiated by methylation and completed by antibiotic binding. ***A***, The tip of H69 in unmethylated *Msm* constituted by A2137 and C2138 is proximal to h44 residues C1392 and A1391. A2137 occupies the capreomycin binding pocket and C2138 is unfavorably positioned with respect to an incoming capreomycin antibiotic, reducing its affinity for the ribosome. ***B***, TlyA-mediated methylation of the 50S results in the transmission of an allosteric signal to the H69 tip, promoting a more open conformation. Within the assembled 70S ribosome, this conformation is preserved by the methyl group of 16S rRNA nucleotide C1392 (shown as a sphere) and the base of A1391. Rotation of A1477 compared to the unmethylated ribosome is also essential to avoid clashes and thereby assists in maintaining the open conformation. ***C***, Upon capreomycin (yellow) binding to the *M. tuberculosis* 70S ribosome (PDB code 5V93), A2151^*Mtb*^ (A2137) flips out of the binding pocket concurrent with a slight rotation of C2152^*Mtb*^ (C2138) in H69. h44 nucleotide A1485^*Mtb*^ (A1476) and A1486^*Mtb*^ (A1477) flip from the antibiotic binding site to accommodate the bound capreomycin. Note that although *Mtb* has active TlyA^II^, the methyl group on Cm1402^*Mtb*^ was not modeled (location is indicated by a black circle).

Allosteric regulation of ribosome function can occur in response to antibiotic binding. For example, macrolide binding at the peptidyl transferase center can cause sensing of leader peptide sequences traversing the nascent peptide exit tunnel and allosterically alter surrounding nucleotides to cause ribosome stalling (43,44). In this mechanism, the leader peptide sequences precede macrolide-resistance genes and stalling is required for their expression. In contrast, it is relatively unknown how RNA modifications *facilitate* antibiotic binding as in the TlyA-mediated case. Recent computational studies on allosteric coupling of different ribosome regions also reveal that H69, particularly 23S rRNA nucleotides A1912^*Eco*^ and U1917^*Eco*^, serves as an allosteric link between the 50S and 30S subunits (45). Building on and consistent with this theme, our analyses suggest that conformational changes beginning at the site of the ribose methylation at C2144 are allosterically transmitted to the H69 tip (C2144 to A2136) which forms part of the capreomycin binding site located at the subunit interface. No other significant conformational changes occur upon methylation of H69, suggesting that this pathway is highly specific. The H69 conformational changes induced by methylation at C2144 are also clinically relevant because *Mtb* clinical isolates containing an A2154^*Mtb*^ deletion within H69 (A2140; A1916^*Eco*^) increase susceptibility to capreomycin (8). The A2140 deletion also results in loss of the C2144 methylation that together, increase the capreomycin MIC by >16-fold in *Msm*. Interestingly, the reduction in susceptibility to capreomycin upon A2140 deletion alone is more pronounced than for loss of TlyA alone, suggesting that restricting the conformational plasticity of H69 impacts capreomycin binding to a larger extent. The importance of A2140 is consistent with helical analyses from our cryo-EM structures and MD simulations in which this nucleotide is consistently adjacent to the sites of major differences between methylated and unmethylated ribosomes (i.e., base steps 8 and 9 within the 8-12 region; **Fig. 3**). Together, these data suggest that A2140 may be a focal point in the allosteric relay of information from the site at C2144 and the tip of H69 that is critical for optimal capreomycin activity.

Although no large conformational changes are induced upon methylation of C1392 on the 30S subunit, there is evidence that this second modification by TlyA^II^ may play an important but supporting role in promoting optimal capreomycin activity. First, in growth competition assays in the presence of capreomycin, an *Eco* strain expressing a TlyA^II^ variant that only partially methylates Cm1409^*Eco*^ (Cm1392) outcompeted a corresponding strain with fully modified C1409^*Eco*^ (11). Second, identification of 16S rRNA mutations in *Mtb* clinical isolates and the finding that corresponding strains of *Msm* (A1391G and C1392U) have capreomycin MICs that are increased by 8-to 16-fold, respectively (16,46) further support the idea that, while clearly less important than Cm2144, Cm1392 does seem to impart cellular fitness advantages important for full capreomycin activity.

Since both the Cm1392 and Cm2144 modifications are catalyzed by TlyA^II^ only in the context of the individual 30S and 50S subunits (11), the Cm1392 modification may help maintain the accessibility of the capreomycin binding pocket in its open conformation during the assembly of the 30S and 50S into a competent 70S complex. Additionally, some local conformational changes in adjacent nucleotides on the complementary strand are observed. Normally, the nucleobases of 16S rRNA A1477 and 23S rRNA A2137 at the tip of the methylated H69 form stacking interactions in the context of the 70S ribosome. Upon C1392 methylation, these two nucleotides would clash but the open conformation of H69 seems to reorganize this interaction to maintain stacking (**Fig. 4C**). This coordinated conformational change thus ensures maintenance of the h44-H69 interaction and, as a result, the open conformation of the H69 tip (35,37). Comparison of the Cm1392 and A1391 positions in the unmethylated and methylated *Msm* ribosomes with the corresponding residues in *Mtb* 70S shows considerable conservation of their interactions in each structure (**Fig. 5**). This structural conservation contrasts with the process of the pocket formation by H69, where methylation and subsequently capreomycin binding affect both the residues involved in forming the binding pocket. We thus propose that H69 methylation generates the open conformation of the capreomycin binding pocket, and h44 methylation ensures that this conformation remains open after the 50S and 30S subunits assemble to form the 70S. Finally, the nature of the capreomycin-bound state of h44 can be observed from the structure of the *Mtb* 70S ribosome-capreomycin complex, in which the residues A1485^*Mtb*^ (A1476) and A1486^*Mtb*^ (A1477) flip from the h44 RNA helix, in addition to the displacement of H69 nucleotide A2151^*Mtb*^ (A2137) discussed above, to accommodate and interact with capreomycin (**Fig. 5C, Supplementary Fig. S6C,F** and **Supplementary Movie 1**).

Overall, our findings elucidate the necessity of a distant ribose methylation for capreomycin binding and highlight the importance of a previously unappreciated pathway of allosteric signaling within 23S rRNA H69. Although capreomycin has a long history of use as an anti-TB agent, it has recently been excluded by the WHO due to its poor performance compared to other current treatments (47). However, efforts are underway to explore combination therapies of capreomycin with other drugs (48), and future targeting of the tuberactinomycin binding site with new generations of antimicrobials may prove fruitful in developing much needed treatments for multi-drug-resistant TB. The role of distant methylation-mediated changes in antibiotic binding sites such as those revealed here may thus represent an important new consideration for designing more effective small molecules targeting the bacterial ribosome.

## DATA AVAILABILITY

Structural coordinates for the TlyA-methylated and unmethylated 70S ribosomes have been deposited in the Protein Data Bank (PDB) with accession codes 9E0P and 9E0N, respectively. Corresponding cryo-EM maps have been deposited in the Electron Microscopy Data Bank (EMDB): 47365 and 47363.

## SUPPLEMENTARY DATA

Supplementary Data are available at NAR online.

## ACKNOWLEDGEMENTS

We thank Dr. James Posey and colleagues at the Centers for Disease Control and Prevention, Atlanta, GA, for providing *M. smegmatis* strains (LR222 and LR222 C101A), and Dr. Ravi K. Koripella for assistance in specimen preparation for cryo-EM.

