## Supplemental Materials for "Distant Ribose 2’-O-Methylation of 23S rRNA Helix 69 Pre-Orders the Capreomycin Drug Binding Pocket at the Ribosome Subunit Interface"

<u>This file contains</u>:

Supplementary Figures S1-S9

Supplementary Tables S1-S2

Still image and Legend for Movie S1

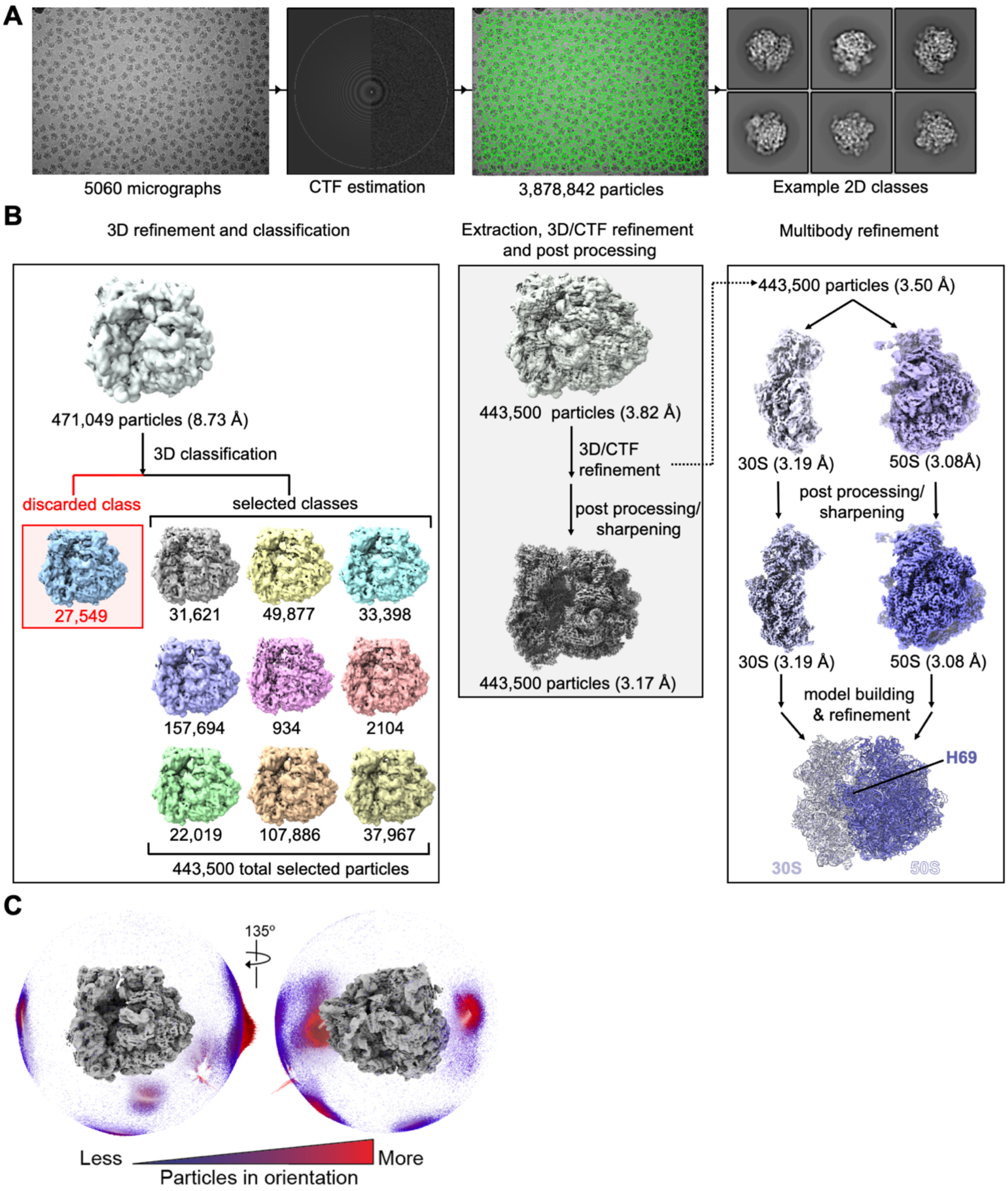


**Fig S1. Workflow for cryo-EM structure determination of the TlyA-methylated *M. smegmatis* (*Msm*) 70S ribosome.** ***A***, Sample micrograph, CTF estimation, particle picking, and example 2D classes. ***B***, The 3D refinement workflow resulting in the final 70S ribosome structure. Initial 3D refinement produced a map at 8.73 Å resolution with a 100 Å box size (upper left), and was followed by a 3D classification to separate particles with poor map quality (lower left). Following rescaling to 400 Å (upper center), multiple rounds of 3D/CTF refinement were used to obtain the final map after postprocessing (lower center). The map obtained prior to postprocessing was segregated into the 50S and 30S subunits, which were subjected to multibody refinement to improve the map resolution (upper right). After postprocessing, the resulting maps were used for both individual model building and refinement in Phenix, and the final models merged to produce the structure depicted in the lower right. ***C***, Angular distribution plot showing the contribution of particles to the final map (from 443,500 particles after 3D/CTF refinement). The left view coincides with that displayed in previous figures. The height of the cylinder bars in the plot (low to high) and their color (blue to red) indicate the number of particles in each view.

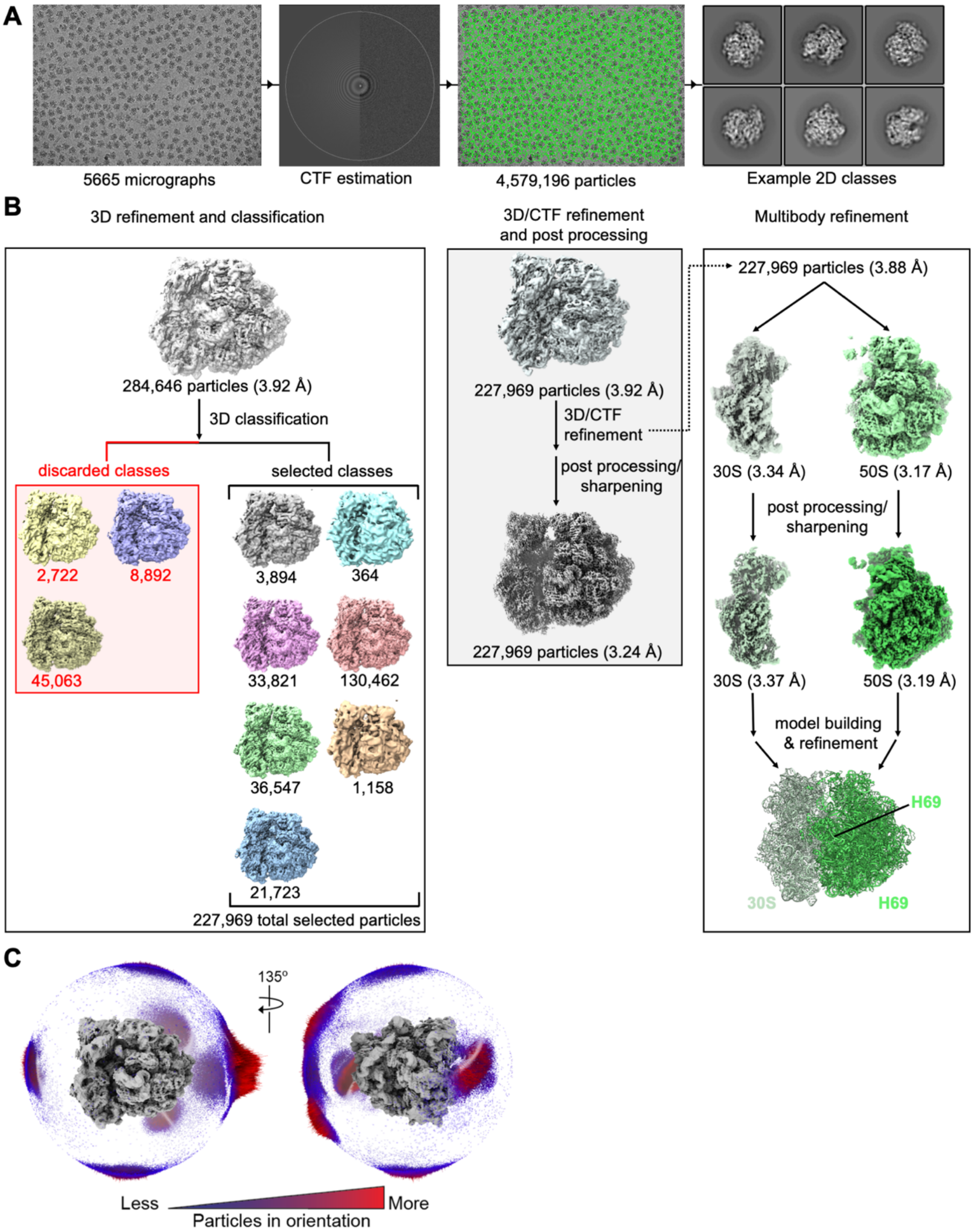


**Fig S2. Workflow for cryo-EM structure determination of the TlyA-unmethylated *Msm* 70S ribosome.** ***A***, Sample micrograph, CTF estimation, particle picking, and representative 2D classes. ***B***, The 3D refinement workflow resulting in the final 70S ribosome structure. Initial 3D refinement resulted in a map at 3.92 Å (upper left) and 3D classification was carried out to distinguish particles with low map quality (lower left). After several rounds of refinement, which included multiple 3D/CTF refinement iterations, the final map was obtained through postprocessing (lower center). The map obtained prior to postprocessing was segregated into the 50S and 30S subunits, which were subjected to multibody refinement to improve the map resolution (upper right). After postprocessing, the resulting maps were used for both individual model building and refinement in Phenix, and the final models merged to produce the structure depicted in the lower right. ***C***, Angular distribution plot showing the contribution of particles to the final map (from 227,969 particles after 3D/CTF refinement). The left view coincides with that displayed in previous figures. The height of the cylinder bars in the plot (low to high) and their color (blue to red) indicate the number of particles in each view.

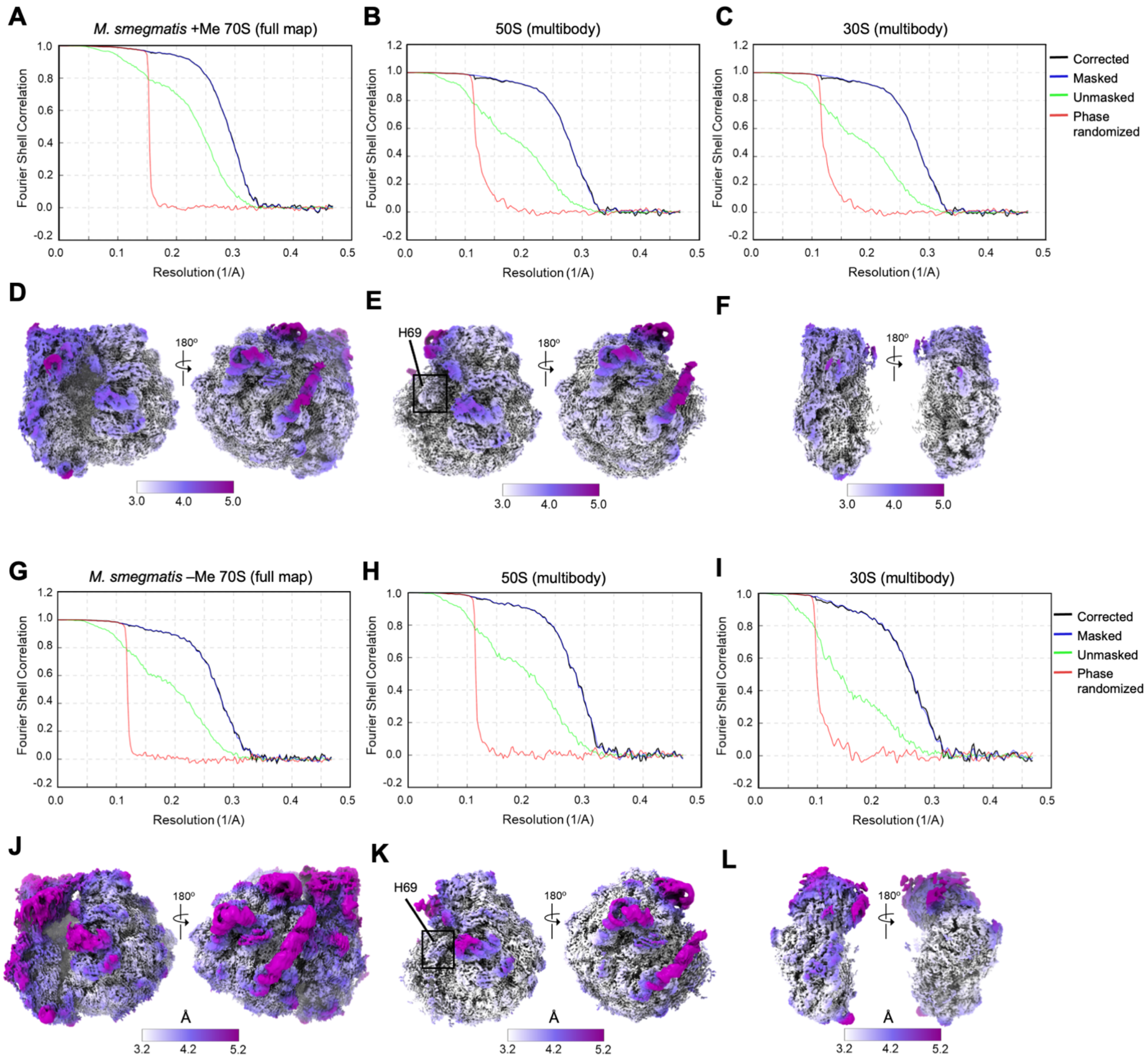


**Fig S3. Analysis of cryo-EM map resolution.** Fourier shell correlation (FSC) curves for ***A***, the full *Msm* TlyA-methylated 70S map, and methylated ***B***, 50S and ***C***, 30S subunit maps from multibody refinement. Curves shown are: corrected (black; under blue line where not visible), masked data (blue), unmasked data (green), and phase randomized data (red). Maps colored to show local resolution of ***D***, the full TlyA-methylated 70S map, ***E***, 50S subunit, and ***F***, 30S subunit. The resolution of the map was determined from the 0.143 FSC intercept of the corrected data, and values are indicated in the scale bar. ***G-L***, as for panels A-F but for the TlyA-unmethylated ribosome.

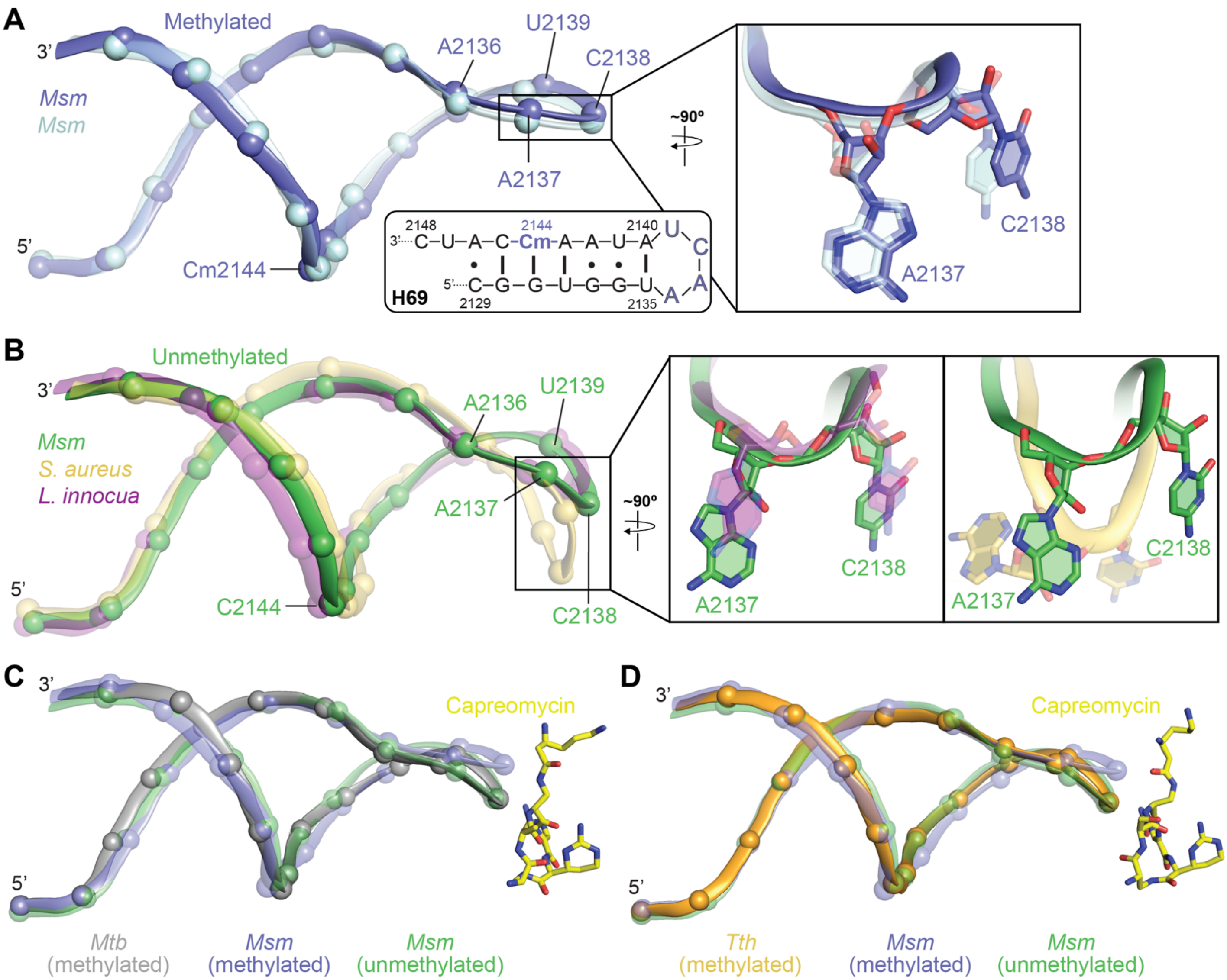


**Fig S4. Superimposition of methylated and unmethylated H69 from different bacterial species showing the differences at the tip of the helix.** ***A***, Methylated H69 from a previously published structure of *M. smegmatis* ribosomes (PDB: 5O61; light blue) overlays well with our present structure (slate), indicating their similarity. *Inset,* superimposition of the nucleotides (A2137 and C2138) at the H69 tip show that they are positioned identically. ***B***, Unmethylated H69 from *Msm* (green) superimposes well with unmethylated H69 of *L. innocua* (PDB: 8UU4, purple) and *S. aureus* (PDB: 7TTU, gold) ribosomes, thereby reflecting their closeness. *Inset,* superimposition of the nucleotides (A2137 and C2138) at the H69 tip from *Msm* and *L. innocua* show that they are positioned similarly. However, the nucleotides at the H69 tip from *S. aureus* are shifted further compared to the nucleotides in *Msm* and *L. innocua*. ***C***,***D***, H69 of *Mtb* (PDB: 5V93; gray) and *Tth* ribosomes (PDB: 4V7M; orange) with bound capreomycin (yellow) exhibits a position similar to the unmethylated H69 (light blue) of *Msm*, although the nucleotides at the tip are oriented differently as noted previously.

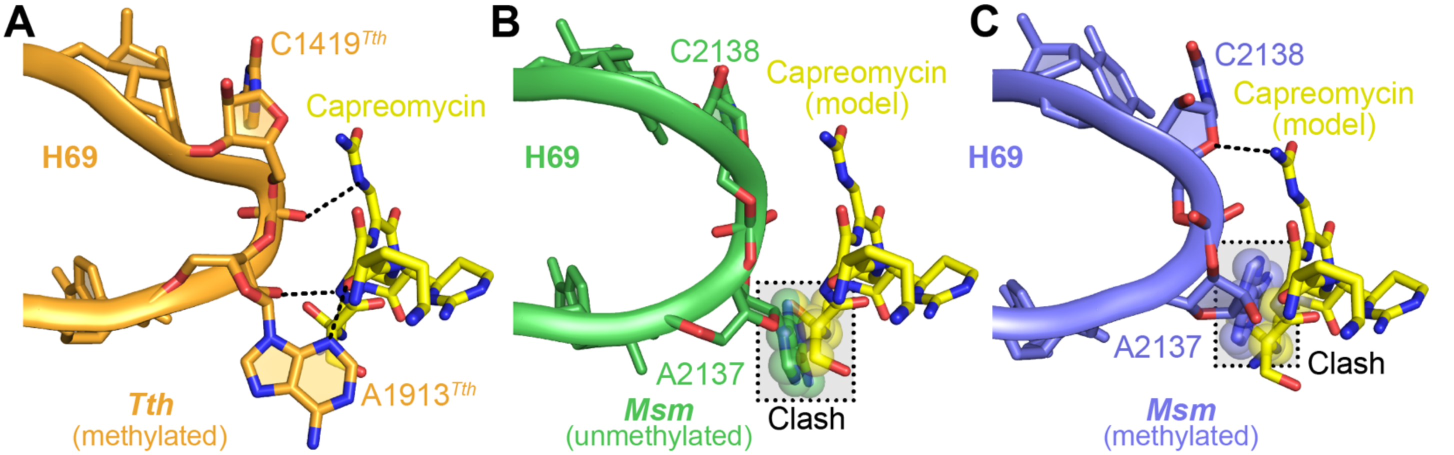


**Fig S5. H69 nucleotides at the tip of methylated and unmethylated *Msm* 70S undergo unique changes compared to *Tth*.** ***A***, Capreomycin (yellow) is stabilized in the antibiotic binding pocket via hydrogen bonds with residues A1913$^{Tth}$ and C1914$^{Tth}$. ***B***, In the unmodified *Msm* ribosome, A2137 obstructs the binding site and would clash with capreomycin (yellow; shown modeled in the *Msm* structure). C2138 shows no potential interaction with the modeled capreomycin. ***C***, In the unmodified *Msm* ribosome, C2138 and the end of the H69 pivot away from the binding pocket, partially opening it up for capreomycin entry and binding. This movement may enable initial contact between the nucleotide and the antibiotic. The clashing atoms of A2137 and capreomycin are shown as semi-transparent spheres. Dashed lines (black) indicate hydrogen bonds.

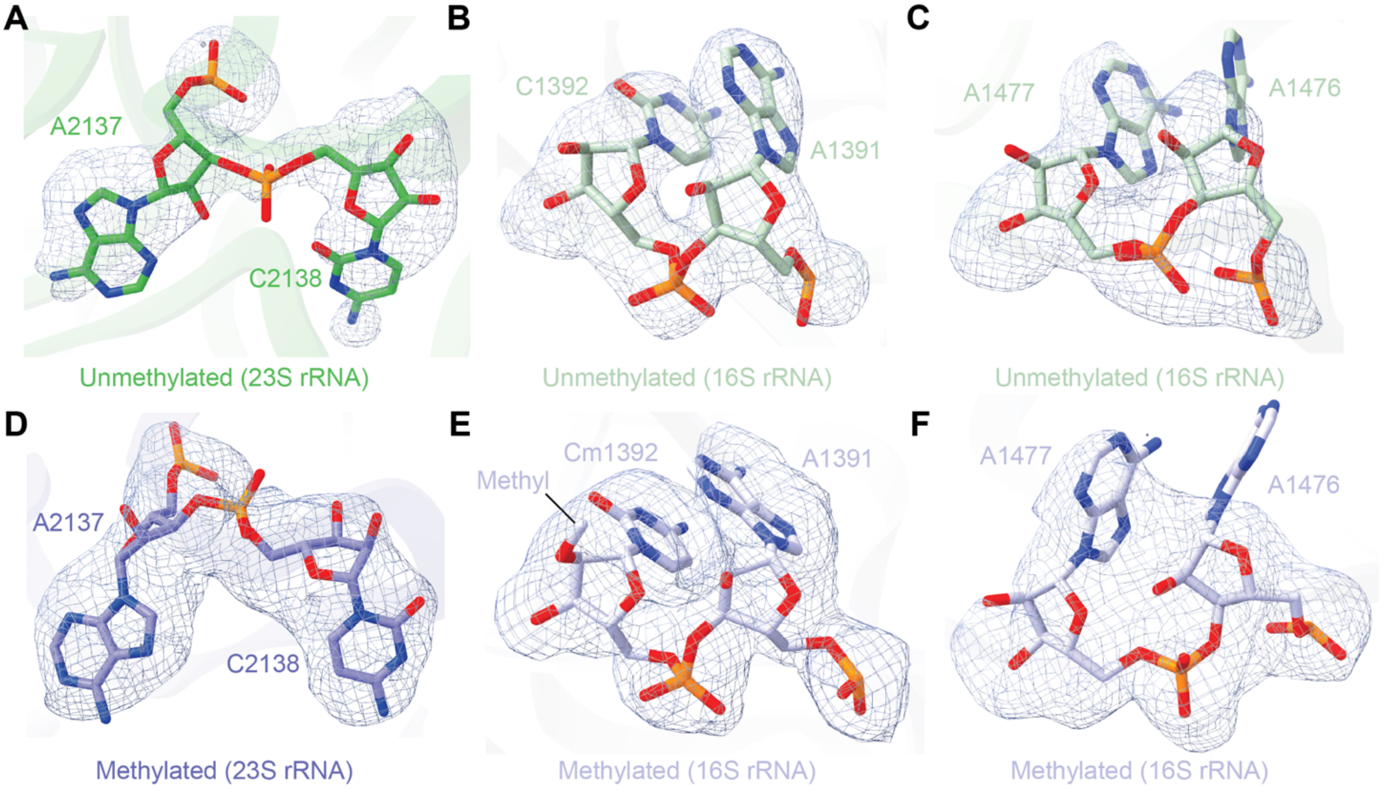


**Fig S6. Cryo-EM map quality of nucleotides in the TlyA-methylated and unmethylated *Msm ribosomes*.** The nucleotides discussed in the main text are shown with the final complete map (blue mesh) from both 70S ribosomes. The residues are colored as in main text figures. All the maps are obtained from DeepEMhancer sharpening of the multibody maps. Unmethylated *Msm* ribosome residues are shown in ***A***, 2137 and 2138, ***B***, A1391 and C1392, ***C***, A1476 and A1477; and methylated *Msm* ribosome residues are shown in ***D***, A2137 and C2138, ***E***, A1391 and C1392, and ***F***, A1476 and A1477.

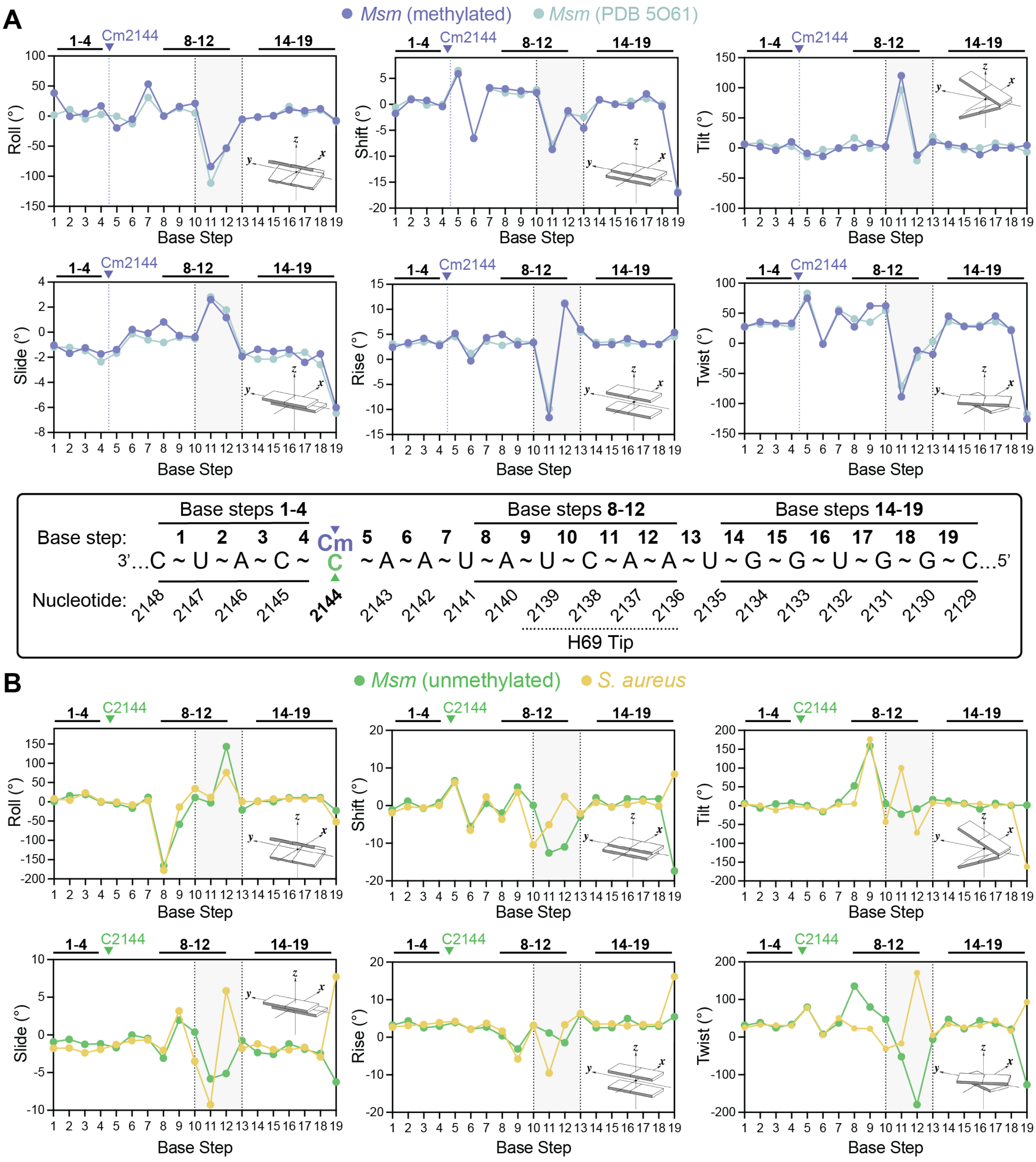


**Fig S7. H69 helical parameters are similar between independent TlyA-methylated *M. smegmatis* structures (both) and unmethylated structures from *M. smegmatis* and *S. aureus. A***, All six helical parameters assessed (roll, shift, twist, slide, rise, and shift) are similar between the methylated H69s from the *Msm* structure described in the present study and a previously reported structure (PDB 5O61). Also indicated on the plot and schematic of the H69 sequence are the positions of each base step and site of methylation (Cm2144). The three regions of base steps indicated (1-4, 8-12, and 14-19) are those discussed in the main text. ***B***, Comparison of the same helical parameters between for unmethylated H69 from *Msm* (present work) and *S. aureus*; dissimilarities are confined mostly at the tip (gray shaded region), which is expected as the nucleotides at the tip of *S. aureus* adopts a unique conformation compared to *Msm.*

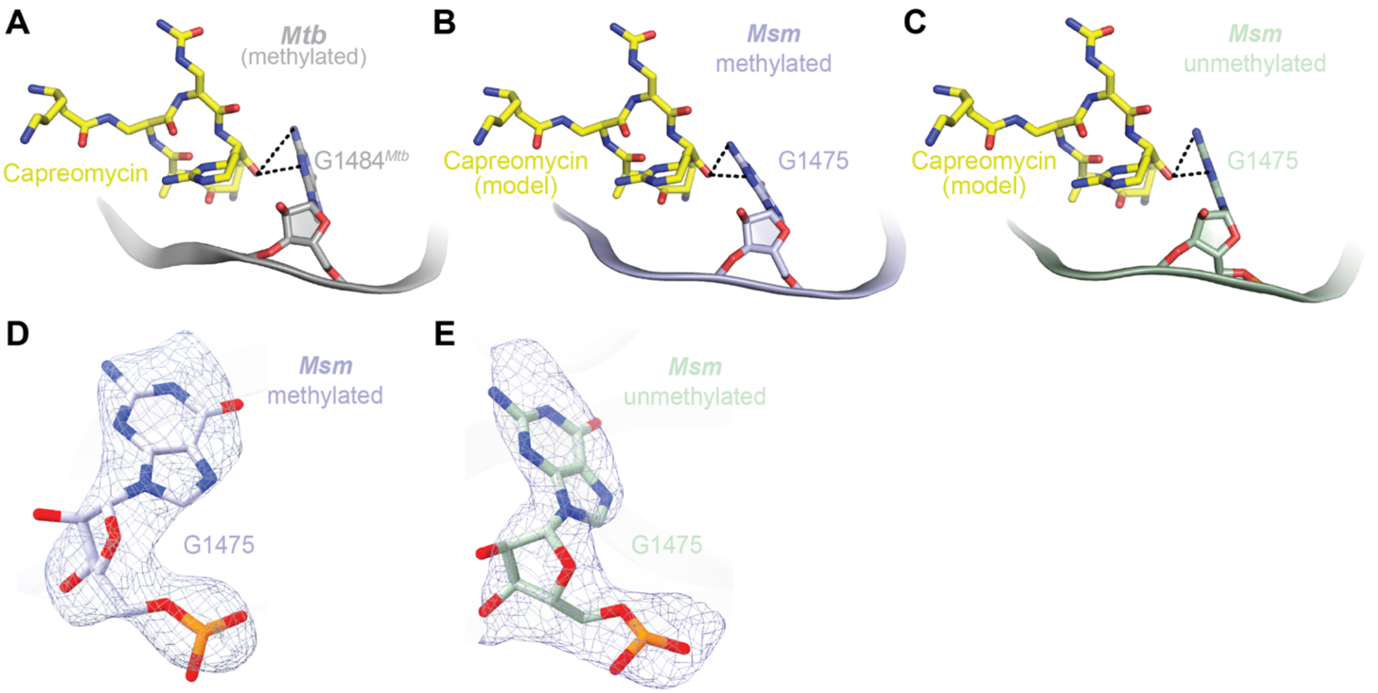


**Fig S8. The position of G1475 in h44 is not affected by methylation status.** ***A***, G1484$^{Mtb}$ in methylated *Mtb* (PDB 5V93; gray) anchors the capreomycin (yellow) in the binding pocket. Similarly, irrespective of methylation status, G1475 in ***B***, methylated (light blue) or ***C***, unmethylated (light green) *Msm* is observed to be within hydrogen bonding distance (dashed black lines) with modeled capreomycin (yellow), showing that the position of the nucleotide is independent of the modification. The modeled G1475 in ***D***, methylated and ***E***, unmethylated *Msm* ribosome structure are shown with the surrounding maps.

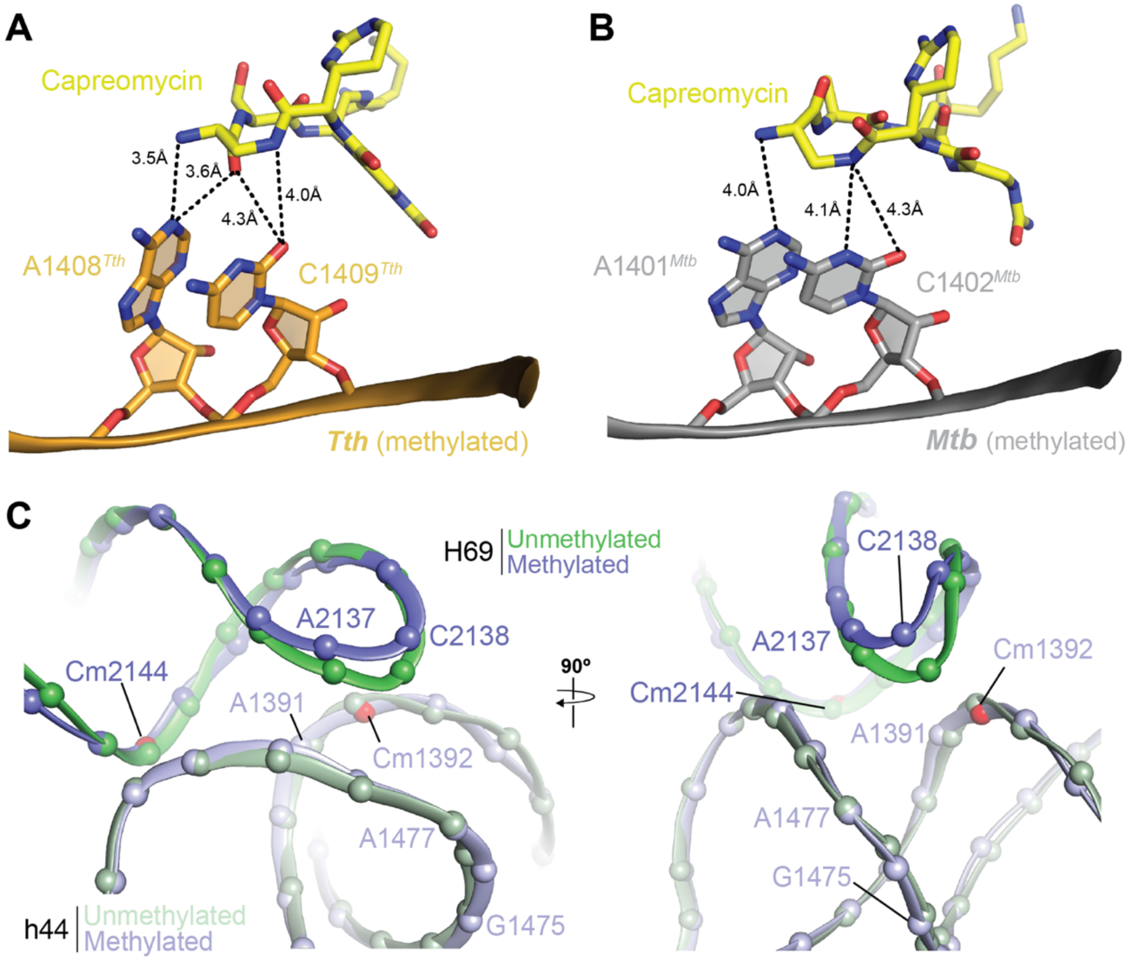


**Fig S9. The capreomycin binding pocket is formed by H69 of the 50S and h44 of the 30S.** ***A***, A1408$^{Tth}$ and C1409$^{Tth}$ in *Tth* (PDB 4V7M; orange) and their equivalent nucleotides in ***B***, *Mtb* A1401$^{Mtb}$ and C1402$^{Mtb}$ (PDB 5V93; gray) have been proposed to form long-range hydrogen bonding interactions (dashed black lines) with capreomycin (yellow) to stabilize the antibiotic in the binding pocket. ***C***, Superimposition of the methylated and unmethylated H69 and part of h44 of *Msm* with key nucleotides involved in either formation of the capreomycin binding pocket or direct antibiotic binding labeled.

### SUPPLEMENTAL TABLES

**Table S1. RMSD for structural alignment of H69 from methylated and unmethylated *Msm* ribosomes and those from different organisms.**

| *Msm* 70S structure (present study) | Species | H69 methylated | PDB | RMSD | Atoms |
|---|---|---|---|---|---|
| Methylated (+TlyA) PDB 9E0P | *Msm* | + | 5O61 | 0.86 | 393 |
| | *S. aureus* | - | 7TTU | 1.01 | 325 |
| | *L. innocua* | - | 8UU4 | 0.89 | 373 |
| | *Msm* | - | 9E0N | 1.10 | 382 |
| | *M. tuberculosis* | + | 5V93 | 1.07 | 389 |
| | *T. thermophilus* | + | 4V7M | 0.95 | 367 |
| Unmethylated (-TlyA) PDB 9E0N | *M. tuberculosis* | + | 5V93 | 1.13 | 388 |
| | *T. thermophilus* | + | 4V7M | 0.82 | 342 |

**Table S2. Standard deviation calculation of the difference between the helical parameters of methylated and unmethylated H69.**

| Helical parameter | Average difference standard deviation[a] | | |
|---|---|---|---|
| | Base steps 1-4 | Base steps 5-9 | Base steps 14-19 |
| Roll | 11 | 63 | 6.0 |
| Shift | 0.6 | 1.7 | 0.9 |
| Twist | 3.0 | 43 | 6.0 |
| Slide | 0.5 | 1.6 | 0.4 |
| Rise | 0.6 | 2.0 | 0.3 |
| Tilt | 4.0 | 63 | 3.0 |

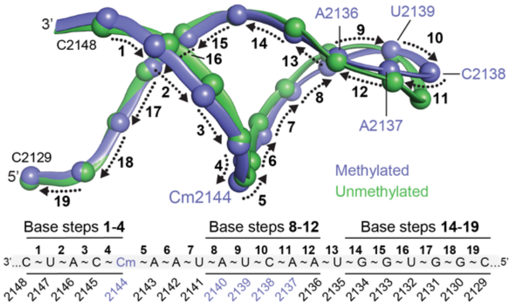


[a]Helical parameter differences between methylated and unmethylated H69 were calculated for each base step and the values averaged for base steps within each of the three H69 regions. The standard deviation of these average calculations is consistently highest for the region spanning from near the site of methylation to the first nucleotide at the tip of H69 (base steps 5-9).

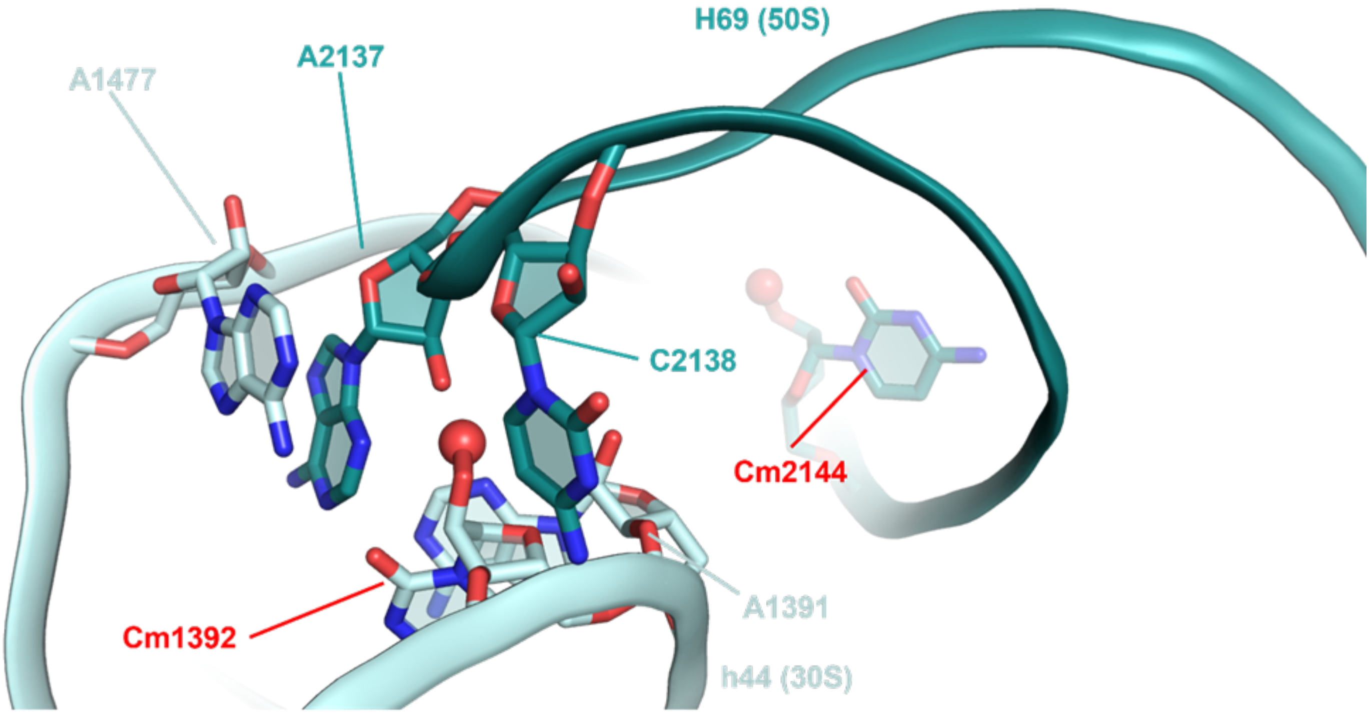


**Movie S1. Still image from the movie depicting the compact-to-open conformational transition of the capreomycin binding pocket.** The addition of a methyl group (red sphere) on C2144 (teal) initiates the propagation of structural changes from the site of methylation to the tip of H69 in the 50S subunit (residues A2137 and C2138). The methyl group (red sphere) of C1392, the base of A1391, and A1477 in h44 of the 30S (pale cyan) preserves the open conformation of H69 tip by partially occupying the locations of H69 nucleotides A2137 and C2138 (teal) in the unmethylated ribosome and therefore potentially causing steric clashes if the two nucleotides fail to maintain their methylated conformation. However, capreomycin (yellow) binding is prevented in the open conformation due to steric clashes with A2137 and A1477, which are relieved as both the residues flip out of the binding pocket. The synergistic effect of the two methylations and finally capreomycin binding opens the antibiotic binding pocket and primes it for drug binding.
